# A covalent irreversible inhibitor binds in two mutually exclusive conformations to the active-site cysteine residue of human aldehyde dehydrogenase 1A3

**DOI:** 10.64898/2026.07.14.738401

**Authors:** Daniela Covaleda, David Vizarraga, Tulsi Upadhyay, Jiyun Zhu, Daniel Abegg, Raquel Pequerul, Martín Hugo, Alexander Adibekian, Ignacio Fita, Xavier Parés, Francesc Xavier Avilés, Matthew Bogyo, Jaume Farrés

## Abstract

Aldehyde dehydrogenases (ALDH) are enzymes that catalyze the NAD(P)^+^-dependent oxidation of aldehydes into carboxylic acids, playing roles in detoxification, biosynthesis, and regulatory functions. Dysfunction of ALDH is associated with serious conditions such as alcohol intolerance, cancer, cardiovascular problems, and neurological disorders. In humans, ALDH1A1 and ALDH1A3 isoforms act as retinaldehyde dehydrogenases and are overexpressed in various cancers, where high levels are associated with increased tumor malignancy, cancer stem cell traits, and therapeutic resistance. ALDH1A3 is recognized as a promising target for anticancer therapies, with several inhibitors, mainly reversible, developed to specifically target it or the enzyme family.

Since ALDH enzymes can also display esterase activity, we used this property to develop an *in vitro* assay specifically targeting the esterase function of ALDH1A3. A highly conserved active-site cysteine in ALDH1A3 is located at the bottom of two converging channels, which define the substrate- and cofactor-binding pockets. To target this catalytic cysteine, we screened a library of 3,200 cysteine-focused covalent fragments. This led to the identification of Z3405279217 (Z34), an acrylamide-based covalent compound that inhibits both ALDH1A1 and ALDH1A3 at sub-micromolar levels. Biochemical and biophysical tests confirmed that Z34 acts as a time-dependent, covalent, and irreversible binder to the active-site cysteine. In this work, we determined the Cryo-EM structure of the ALDH1A3-Z34 complex at 2.26 Å resolution, confirming the covalent attachment to the catalytic cysteine of Z34. Notably, two mutually exclusive covalent binding modes were observed: one occupying the substrate-binding pocket and the other the cofactor-binding region. Z34 displayed unexpected binding modes within the active site and holds promise as a lead compound for future drug development.

**GRAPHICAL ABSTRACT:** 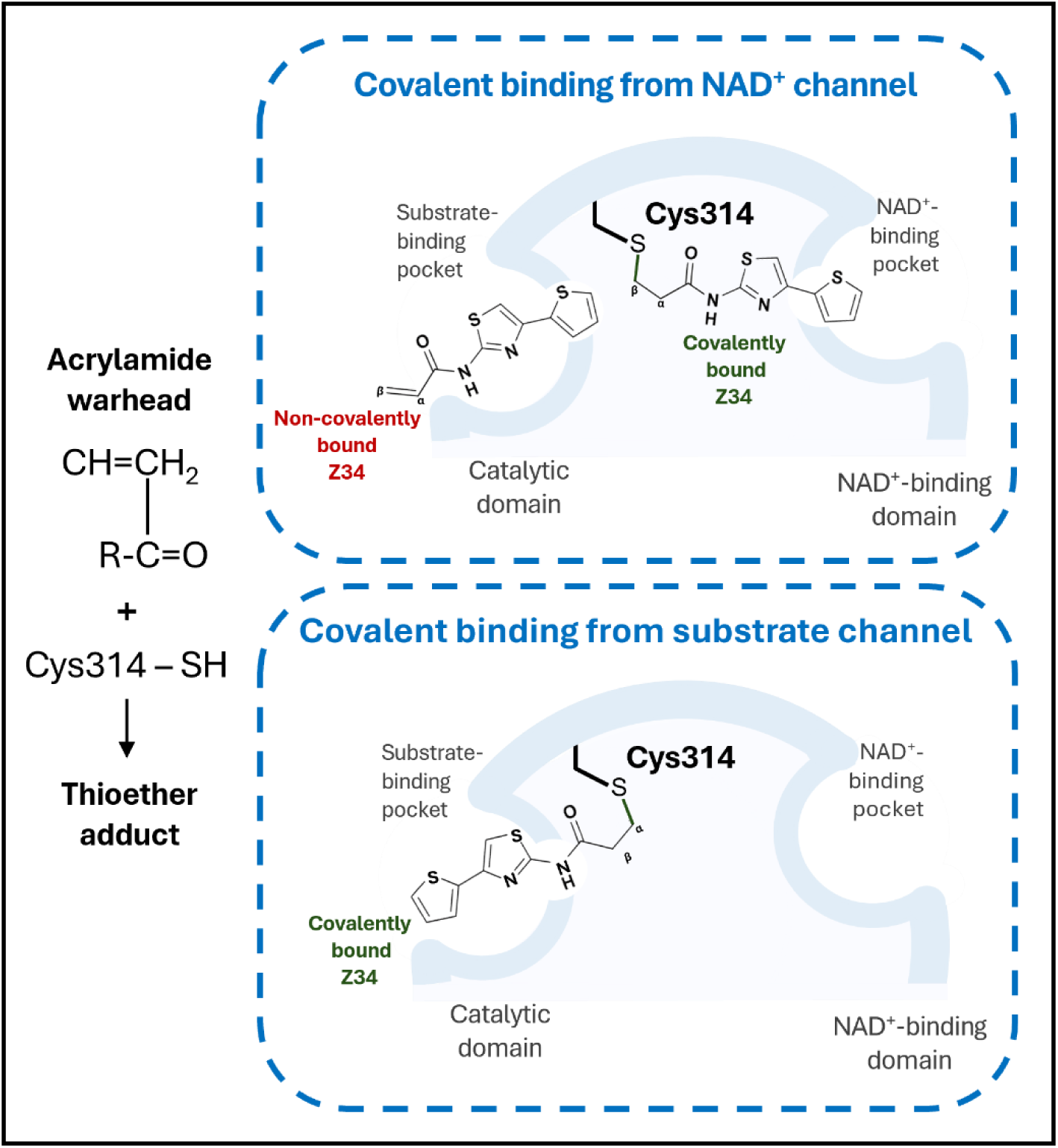

## INTRODUCTION

Aldehyde dehydrogenase (ALDH) catalyzes the NAD(P)^+^-dependent oxidation of aldehydes to their corresponding carboxylic acids and plays a central role in cellular detoxification and metabolic regulation. Among the human isoforms, ALDH1A1 and ALDH1A3 are retinaldehyde dehydrogenases and are frequently upregulated in a wide range of cancers, where they are associated with tumor aggressiveness, cancer stem cell phenotypes, and resistance to therapy (1–3). Several reports have demonstrated that ALDH1A3 is a validated target for anticancer therapy (1,2,4–8).

ALDH1A isoforms are homotetrameric enzymes composed of four subunits of 55–57 kDa. Each monomer contains catalytic, cofactor-binding and oligomerization domains. A strictly conserved active-site cysteine residue (Cys314 in ALDH1A3) lies at the bottom of two converging channels which define the substrate- and cofactor-binding pockets. ALDH catalytic mechanism relies on the nucleophilic attack of the substrate carbonyl carbon by the thiol group of Cys314, yielding a thiohemiacetal intermediate followed by deacylation and hydride transfer to NAD^+^ (9). Interestingly, beyond their aldehyde dehydrogenase function, ALDH enzymes also exhibit cofactor-independent esterase activity, hydrolyzing ester substrates into the corresponding carboxylic acids and alcohols (10–12). This hydrolytic capability is mechanistically related to that observed in thiol proteases and serine hydrolases. Ester substrates interact with the same catalytic pocket used for aldehyde oxidation in ALDH, and the reaction involves nucleophilic attack by the catalytic Cys residue on the ester carbonyl carbon, similarly to the canonical ALDH catalytic mechanism (13). Recently, we reported the use of fluorescent resorufin-derived esters as substrates of different ALDH isoforms (14).

Several ALDH1A3 inhibitors with moderate selectivity have been developed. Although preferentially targeting ALDH1A3, they may also inhibit other ALDH1A isoforms. Most existing ALDH1A3 inhibitors are reversible with a competitive binding pattern and only few three-dimensional structures are available so far (Table 1). These inhibitors have proven effective against a variety of cancer types, such as glioblastoma, breast, ovarian and prostate cancer cell lines.

**Table 1.**
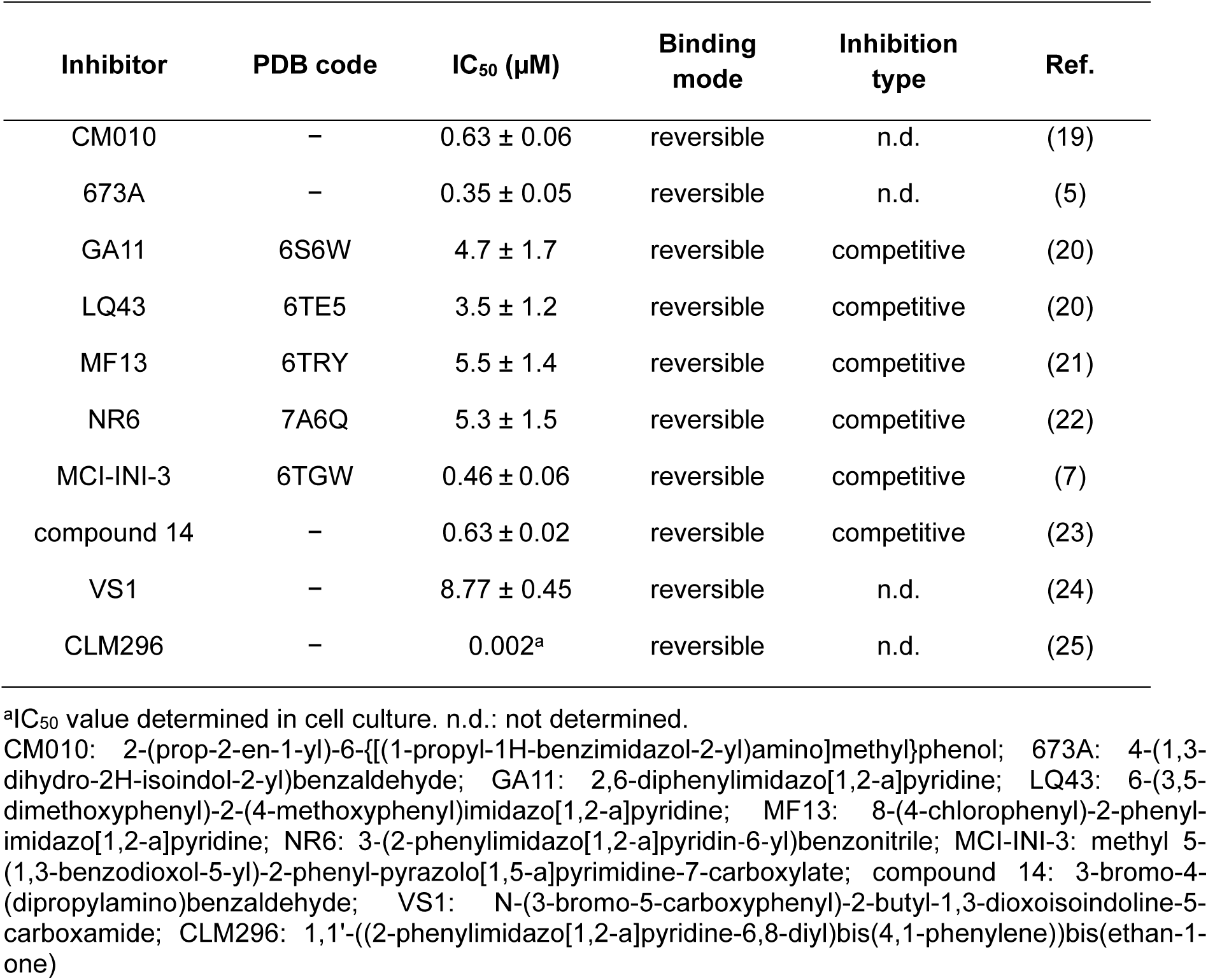
Reversible inhibitor compounds against the ALDH1A3 isoform.

| Inhibitor | PDB code | IC<sub>50</sub> (μM) | Binding mode | Inhibition type | Ref. |
| --- | --- | --- | --- | --- | --- |
| CM010 | – | 0.63 ± 0.06 | reversible | n.d. | (19) |
| 673A | – | 0.35 ± 0.05 | reversible | n.d. | (5) |
| GA11 | 6S6W | 4.7 ± 1.7 | reversible | competitive | (20) |
| LQ43 | 6TE5 | 3.5 ± 1.2 | reversible | competitive | (20) |
| MF13 | 6TRY | 5.5 ± 1.4 | reversible | competitive | (21) |
| NR6 | 7A6Q | 5.3 ± 1.5 | reversible | competitive | (22) |
| MCI-INI-3 | 6TGW | 0.46 ± 0.06 | reversible | competitive | (7) |
| compound 14 | – | 0.63 ± 0.02 | reversible | competitive | (23) |
| VS1 | – | 8.77 ± 0.45 | reversible | n.d. | (24) |
| CLM296 | – | 0.002<sup>a</sup> | reversible | n.d. | (25) |

<sup>a</sup>IC<sub>50</sub> value determined in cell culture. n.d.: not determined.
CM010: 2-(prop-2-en-1-yl)-6-[[[(1-propyl-1H-benzimidazol-2-yl)amino]methyl]phenol; 673A: 4-(1,3-dihydro-2H-isindol-2-yl)benzaldehyde; GA11: 2,6-diphenylimidazo[1,2-a]pyridine; LQ43: 6-(3,5-dimethoxyphenyl)-2-(4-methoxyphenyl)imidazo[1,2-a]pyridine; MF13: 8-(4-chlorophenyl)-2-phenylimidazo[1,2-a]pyridine; NR6: 3-(2-phenylimidazo[1,2-a]pyridin-6-yl)benzonitrile; MCI-INI-3: methyl 5-(1,3-benzodioxol-5-yl)-2-phenyl-pyrazolo[1,5-a]pyrimidine-7-carboxylate; compound 14: 3-bromo-4-(dipropylamino)benzaldehyde; VS1: N-(3-bromo-5-carboxyphenyl)-2-butyl-1,3-dioxoisindoline-5-carboxamide; CLM296: 1,1'-((2-phenylimidazo[1,2-a]pyridine-6,8-diyl)bis(4,1-phenylene))bis(ethan-1-one)

Lately, covalent inhibitors are attracting growing interest in drug discovery, with nearly 30% of recently approved drugs acting through covalent mechanisms (15). Compared with reversible ligands, covalent inhibitors offer several advantages as therapeutic agents and chemical probes. In particular, they can be designed to achieve high selectivity and enhanced potency through sustained inhibition of the intended target. Their prolonged mode of action may also allow reduced dosing and decrease the likelihood of resistance development associated with time-dependent inhibition (16–18). This fact and the lack of covalent ALDH1A3 inhibitors underscores the need for novel strategies to identify covalent binders capable of targeting ALDH1A3.

Here, we describe a high-throughput screening of a cysteine-targeted covalent fragment library of 3,200 commercially available compounds, containing diverse electrophile fragments with preferential reactivity for cysteine nucleophiles. The esterase activity of ALDH was used as a surrogate assay which had been employed successfully in high-throughput assays to screen for ALDH1A1 and ALDH1A3 inhibitors (26–28). Esterase assays do not require the addition of cofactor, reducing assay complexity, cost, and potential interference from compounds affecting cofactor binding or redox cycling. We used a fluorogenic lipid ester, 4-methylumbelliferyl caprylate (4-MUC), as an ALDH substrate which had been previously used for serine hydrolases (29). The screening assay employed a highly sensitive, small-volume format compatible with high-throughput applications, substantially minimizing the consumption of assay reagents and inhibitor compounds. Primary screening identified several compounds, including multiple electrophile classes that had not previously been reported to target the active site of ALDH. Compounds showing low cross-reactivity with cysteine proteases were further selected. Finally, compound Z3405279217 (hereafter referred to as Z34) was used for subsequent biochemical and biophysical characterization, emerging as a time-dependent, irreversible inhibitor. We then determined the first cryo-EM structure of human ALDH1A3 in complex with Z34 at 2.3 Å resolution, directly visualizing covalent modification of the catalytic Cys314 and establishing a structural framework for further structure-based optimization of this initial hit compound.

## RESULTS

### Cysteine-focused covalent fragment library screening against ALDH1A3 esterase activity

To establish a screening platform to identify inhibitors of ALDH1A3 esterase activity, we developed a highly robust 4-methylumbelliferyl (4MU)-based assay of ALDH1A3 esterase activity. For that, we first determined the substrate preference of ALDH1A3 using 4MU substrates with carbon lengths of 2c, 4c, 7c, 8c, and 10c (Figure 1A). We observed that ALDH1A3 strongly prefers substrates with 7c and 8c chain lengths. Its esterase activity for 4-methylumbelliferyl caprylate (8c) was fully inhibited by 400 µM N-ethylmaleimide (NEM), which served as a positive control during high-throughput screening (Figure 1B).

**Figure 1.**
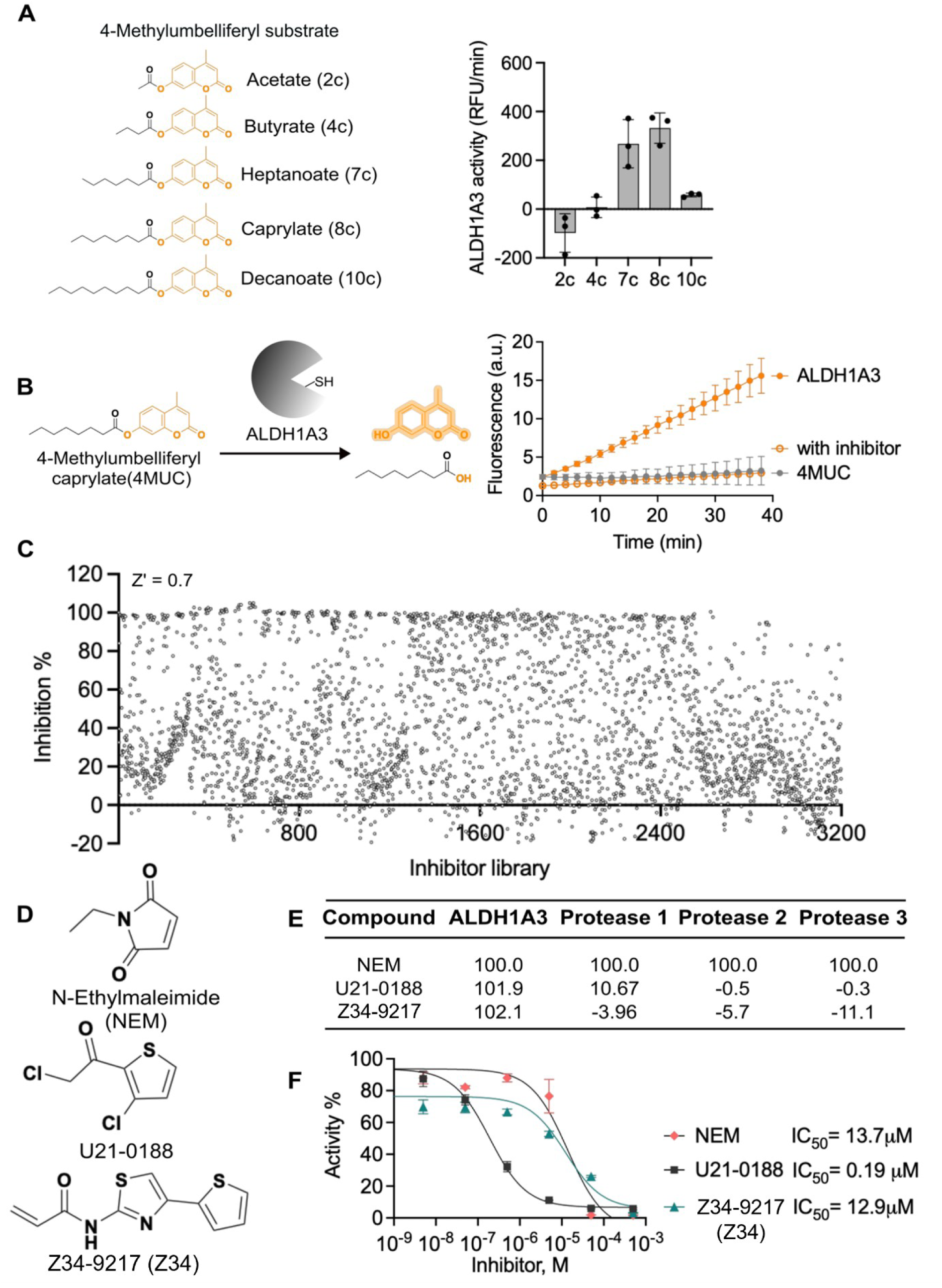
Targeting ALDH1A3 esterase activity. **(A)** Structures of various chain lengths of 4-methylumbelliferyl-based substrates used to evaluate ALDH1A3 substrate preference. The activity plot shows the relative fluorescence units (RFU/min) generated by ALDH1A3 for each compound. Bars represent mean ± SD (n=3). **(B)** Schematic representation of ALDH1A3 esterase activity toward 4-methylumbelliferyl caprylate (4MUC). Representative activity curves for 250 nM ALDH1A3 using 50 μM 4MUC in the absence and presence of 400 μM N-ethylmaleimide (NEM). Values are expressed as mean ± SD of three independent experiments. **(C)** Plot showing the percentage inhibition produced by each compound at 100 μM in the primary fragment library screening against ALDH1A3. NEM was used as a positive control (100% inhibition). **(D)** Chemical structures of NEM and the two selected hits, U21-9018 and Z34-9217 (Z34). **(E)** Comparison of the percentage inhibition of ALDH1A3 and three cysteine proteases by the selected compounds. **(F)** Inhibition curves used to determine the IC_50_ values of each compound against 250 nM ALDH1A3, measured by the initial cleavage rate of 50 μM 4MUC. Percent activity was calculated relative to a DMSO control. Data are represented as mean ± SD (n=3).

We optimized the assay conditions to generate a robust assay with the signal-to-noise ratio or Z’ > 0.7. We used the assay conditions to screen a commercially available 3,200 cysteine-focused covalent fragment library. The library is mainly composed of chloromethyl ketone, chloroacetamide, acrylamide, maleimide, and vinyl sulfone. Because the hit rate was very high, we selected 60 compounds from the initial screening that demonstrated ≍100% inhibition (Figure 1C). To prioritize candidates for further characterization, we wanted to follow only those hits that selectively inhibited ALDH1A3. To assess selectivity, the identified hits were compared with three additional cysteine proteases, and the hit list was narrowed to 40 compounds that specifically inhibited ALD1A3 esterase activity. Further, by dividing the warhead class among the 40 hit compounds, we found 27 maleimide-based hits, 12 chloromethyl ketone-based inhibitors, and only 1 acrylamide-based inhibitor. On the reactivity spectrum, acrylamide shows moderate reactivity in comparison to highly reactive maleimide and chloromethyl ketone. Based on this analysis, to measure inhibitory potency, we selected Z34, the only acrylamide-based hit, and also selected a chloromethyl ketone-based inhibitor, U21-0188. We measured the IC_50_ values for Z34 and U21-0188 and compared them with the broad-spectrum maleimide-based inhibitor NEM. In the inhibition data, U21-0188, a highly reactive electrophile, showed an IC_50_ value of 0.19 ± 0.01 µM. The IC_50_ values for Z34 and NEM were very similar, with IC_50_ values of 12.9 ± 4.5 µM and 13.7 ± 3.2 µM, respectively. To avoid nonspecific reactivity with U21-0188, we proceed with Z34 in the next steps of the hit validation studies.

To assess the structure-activity relationship, we used the DataWarrior tool and generated a 2-D scatter plot of the Structure-Activity Landscape Index (SALI) using OrgFunctions for the 3,200 compounds tested from the library (Figure S1A). In the scatter plot, similar compounds are grouped together, and marker color indicates activity. Further, a closer look at the structure-activity relationship is provided by the neighborhood plots for Z34-9217 (Z34) and U21-0188 (Figure S1B,C). The analysis clearly shows that a small modification to the scaffold of these hit compounds reduces the specificity, as reflected in the percentage inhibition for ALDH1A3 and the other three proteases tested.

### ALDH1A3 dehydrogenase activity inhibition by Z34

Following identification by esterase-based HTS, inhibitor characterization was performed using the hexanal-dependent dehydrogenase activity of ALDH enzymes. Z34 exhibited time-dependent inhibition of both ALDH1A3 and ALDH1A1 (Figure 2A and Figure S4A), a hallmark of covalent or irreversible inhibition, consistent with formation of a stable enzyme–inhibitor complex. Preliminary experiments were performed to establish suitable assay conditions for the biochemical characterization of ALDH1A3 inhibition. Optimal inhibition was observed in 50 mM HEPES, pH 8.0, containing 50 mM MgCl₂, at 25°C, and in the absence of reducing agents. Control experiments assessing the influence of reducing conditions are shown in Figure S3A.

**Figure 2.**
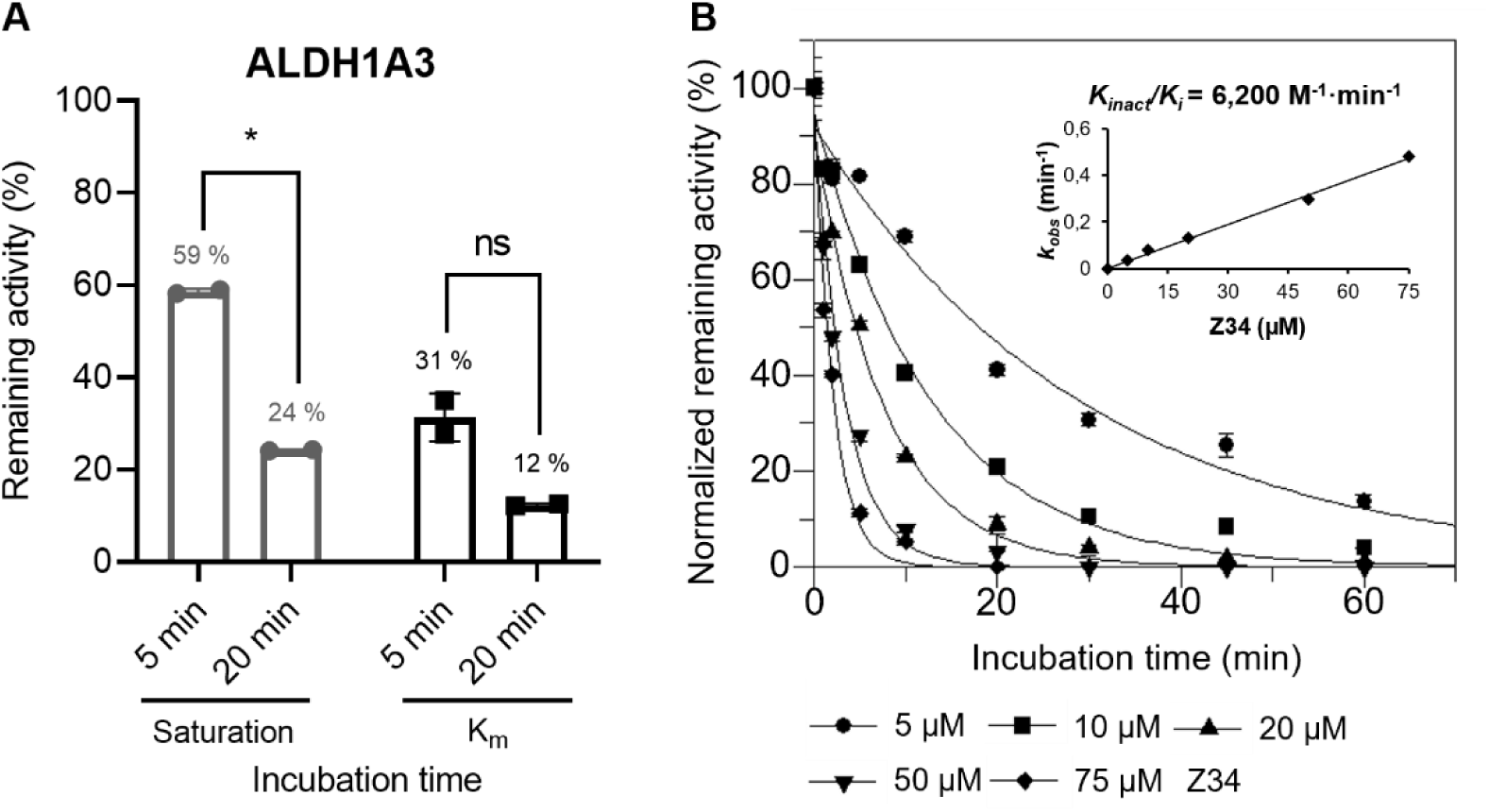
Kinetic characterization of time-dependent ALDH1A3 inhibition by Z34. **(A)** Initial inhibition screening of ALDH1A3 using 10 µM Z34 at saturating (250 µM hexanal) or near-*K_m_* (10 µM hexanal) substrate concentrations after 5- and 20-min pre-incubation. Reactions were performed in 50 mM HEPES buffer containing 50 mM MgCl₂ (pH 8.0) at 25°C. Statistical analysis was performed using one-way ANOVA with multiple comparisons. **(B)** Determination of apparent rate constants (*k_obs_*) for ALDH1A3 at increasing Z34 concentrations (1×, 2×, 4×, 10×, and 20× IC₅₀) using pre-incubation times ranging from 0 to 60 min. The dependence of *k_obs_* on inhibitor concentration was used to calculate *k_inact_*/*K_i_*. Experiments were performed in duplicate, and kinetic parameters were obtained by non-linear regression using GraFit 5.0 (Erithacus Software).

IC₅₀ values determined after 20 min pre-incubation were 5.3 ± 0.6 µM for ALDH1A3 and 2.1 ± 0.1 µM for ALDH1A1 (Figure S2C,D). Apparent inactivation rate constants (*k_obs_*) were measured at increasing inhibitor concentrations (Figure 2B for ALDH1A3; Figure S4B for ALDH1A1). Fitting *k_obs_* as a function of inhibitor concentration yielded *k_inact_* and *K_i_* values, with the ratio *k_inact_*/*K_i_* reflecting inhibitor potency by integrating binding affinity and inactivation rate. For ALDH1A3, Z34 displayed a *k_inact_*/*K_i_* value of 6,200 M⁻¹·min⁻¹, indicating an efficient enzyme inactivation, compared to DIMATE (1,700 M⁻¹·min⁻¹), an ALDH1A inhibitor currently in clinical trials (2). Notably, Z34 exhibited even higher inactivation efficiency toward ALDH1A1 (*k_inact_*/*K_i_* = 11,400 M⁻¹·min⁻¹).

Substrate concentrations for inhibition assays were chosen at saturating hexanal concentrations for each isoform (ALDH1A3: 250 µM, ALDH1A1: 30 µM; Figure S2A,B). Although Z34 also inhibits ALDH1A1, subsequent mechanistic and structural analyses focused on ALDH1A3 to provide a detailed characterization of this isoform, while data for ALDH1A1 are included in the Supplementary Information.

### Competitive effect of NAD⁺ on ALDH1A3-Z34 interaction

To investigate the effect of NAD^+^ on Z34-mediated inhibition of ALDH1A3, enzyme activity was measured in the presence of increasing NAD⁺ concentrations (0–1000 µM). In the absence of NAD^+^, incubation with 75 µM Z34 resulted in complete inhibition of enzymatic activity, whereas increasing NAD⁺ concentrations progressively restored remaining activity, reaching 19.1% at 1000 µM NAD⁺ (Figure 3A). These results indicate that NAD⁺ modulates the initial interaction between Z34 and ALDH1A3; however, the inhibitor retains the ability to inhibit the enzyme even at saturating cofactor concentrations. A similar effect was observed at lower inhibitor concentration (10 µM; Figure S5A), where NAD⁺ also promoted a concentration-dependent recovery of activity.

**Figure 3.**
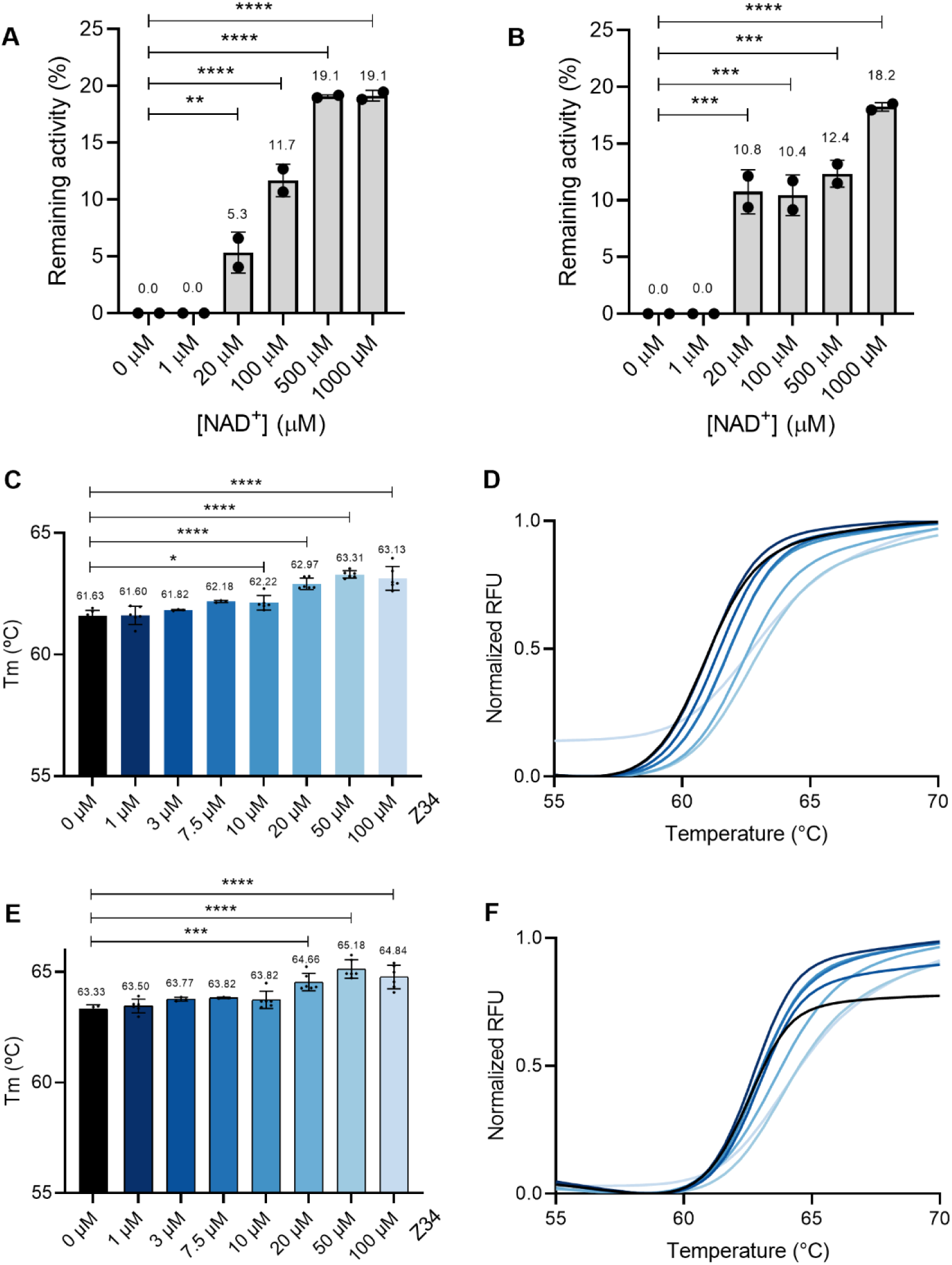
Effect of NAD⁺ on Z34 binding and thermal stabilization of ALDH1A3. **(A)** Bar chart showing ALDH1A3 enzymatic activity after 20 min incubation with 75 µM Z34 in the presence of NAD⁺ at concentrations ranging from 0 to 1000 µM. Enzymatic activity was measured under saturating substrate conditions (250 µM hexanal). Data are presented as mean ± SD of duplicate measurements. **(B)** Bar chart showing ALDH1A3 enzymatic activity after 20 min incubation with 75 µM Z34 in the presence of increasing NAD⁺ concentrations (0–1000 µM). Enzymatic activity was measured at a substrate concentration well below the *K_m_* (2.5 µM hexanal; *K_m_* ≈ 15 µM). Data are presented as mean ± SD of duplicate measurements. **(C)** Bar chart showing the mean melting temperatures (Tm) of 1 µM ALDH1A3 incubated with increasing concentrations of Z34 (1–100 µM) in the absence of NAD⁺. Tm values were derived from Boltzmann fits of normalized melting curves. Protein unfolding was monitored using SYPRO Orange with a temperature ramp from 25°C to 75°C in 0.3°C increments, holding for 1 min at each step. **(D)** Representative sigmoidal melting curves corresponding to the bar chart in panel C. **(E)** Bar chart showing the mean melting temperatures (Tm) of 1 µM ALDH1A3 incubated with Z34 (1–100 µM) in the presence of 500 µM NAD⁺ under the same conditions as in panel C. **(F)** Representative sigmoidal melting curves corresponding to the bar chart in panel E.

A comparable profile was observed under sub-saturating substrate conditions (2.5 µM hexanal; *K_m_* ≈ 15 µM; Figure 3B). At intermediate NAD⁺ concentrations (20–500 µM), remaining activity ranged from 10.4 to 12.4%, increasing to 18.2% at 1000 µM NAD⁺. Overall, the response was consistent with that obtained under saturating substrate conditions, indicating that the extent of inhibition after 20-min pre-incubation is largely independent of substrate concentration. Together, these results support a model in which NAD⁺ modulates initial binding events, whereas time-dependent inactivation leads to a stable inhibited state that is not readily reversed by substrate competition under assay conditions.

To evaluate whether the non-catalytic cysteine adjacent to the catalytic Cys314 contributes to Z34-mediated inhibition, the same experimental conditions were applied to the C313A mutant. A similar behavior was observed compared with wild-type ALDH1A3 (Figure S5B), indicating that Cys313 does not play a major role in Z34-mediated inhibition.

### Effect of NAD⁺ on Z34 binding and thermal stabilization of ALDH1A3

The ALDH1A3-Z34 complex exhibited an increase in melting temperature (Tm) relative to apo ALDH1A3 under both NAD⁺-free (Figure 3C) and NAD⁺-containing conditions (Figure 3E), indicating ligand-induced stabilization of the protein. Notably, the NAD⁺ control displayed higher thermal stability compared to the NAD⁺-free control (ΔTm = 1.7°C), reflecting the intrinsic stabilizing effect of the cofactor. In NAD⁺-free conditions, significant stabilization was observed starting at 10 µM Z34 (ΔTm = 0.6 °C, p < 0.05), with maximal stabilization at 20–100 µM (ΔTm = 1.5°C, p < 0.0001). In the presence of NAD⁺, stabilization became significant at 20 µM (ΔTm = 1.3°C, p < 0.001), and 50–100 µM (ΔTm = 1.5°C, p < 0.0001). Overall, Z34 produced a comparable stabilizing effect on ALDH1A3 in the presence and absence of NAD⁺, suggesting that ligand binding occurs independently of the cofactor and consistently enhances protein stability. Sigmoidal fits of normalized fluorescence curves (Figure 3D,F) were used to determine Tm values, with all measurements performed in duplicate. These results are consistent with the partial inhibition observed in the presence of NAD⁺ (Figure 3A and Figure S5A), confirming that the cofactor hinders but does not prevent Z34 binding.

### Identification of Z34 covalent binding sites in ALDH1A3 by mass spectrometry

Since the ALDH1A3 active site contains the catalytic Cys314 and a neighboring non-catalytic Cys313, ESI-MS analysis of the intact protein was performed to assess Z34-mediated covalent modification.

Incubation of wild-type ALDH1A3 with increasing molar ratios of Z34 (Figure 4A) revealed concentration-dependent covalent modification of the enzyme. The control sample showed a mass increase of 177 Da, consistent with the known α-N-6-phosphogluconoylation of the N-terminal His-tag commonly observed in recombinant proteins expressed in *E. coli* (30,31). Upon 30 min incubation with Z34, additional mass shifts consistent with the molecular weight of the ligand were detected. At a 1:1 Z34-to-monomer ratio (1×), a single covalent adduct corresponding to one Z34 molecule (Δmass = 236 Da) was detected. Increasing the Z34 concentration to 2× and 3× resulted in additional mass shifts corresponding to sequential modification by a second and third Z34 molecule (each Δmass = 236 Da). These results indicate that while a single Z34 modification predominates at equimolar conditions, higher inhibitor concentrations promote additional covalent labeling events.

**Figure 4.**
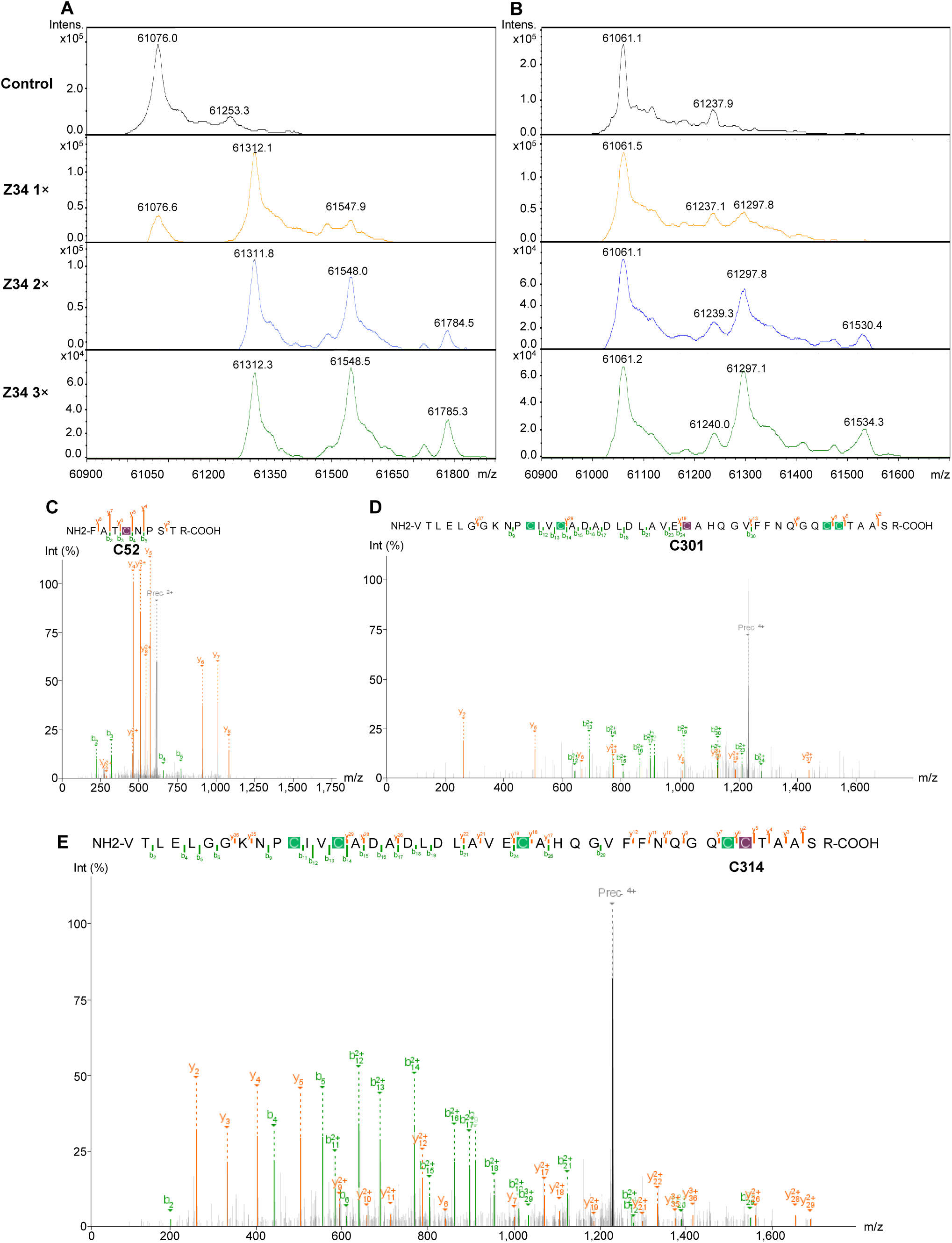
ESI-TOF and MS/MS analysis of ALDH1A3 upon Z34 treatment. **(A, B)** Wild-type ALDH1A3 **(A)** and the catalytic mutant C314S **(B)** were incubated with Z34 at increasing molar ratio per monomer (1×, 2×, and 3×) for 30 min at 25°C. Control samples contained 1% DMSO only. Mass spectra show the monomeric protein species under each condition. Panels are organized from top to bottom as follows: DMSO control, 1× Z34, 2× Z34, and 3× Z34. In the wild-type enzyme, the apo form decreases with increasing Z34 ratio, whereas in the C314S mutant the apo form remains predominant under all tested conditions. Data confirm that Cys314 is selectively modified in the wild-type enzyme. **(C-E)** Identification of labeled cysteine residues by tandem mass spectrometry (MS/MS) following proteolytic digestion. Peptide fragments containing Cys52, Cys301, and Cys314 are shown, demonstrating labeling by Z34. Shown on the sequence in purple is the Z34 adduct and in green iodoacetamide adducts.

Analysis of the catalytic mutant ALDH1A3 C314S (Figure 4B) also revealed the ∼177 Da mass increase associated with the His-tag modification. In contrast to the wild-type protein, the apo form remained the predominant species under all treatment conditions. Under 1× treatment, only a very minor signal consistent with the association of a single Z34 molecule was observed. At 2× treatment, a peak corresponding to one ligand-associated species became detectable, while a weak signal suggestive of a second modification was also present. Under 3× treatment, the apo protein remained the most abundant species, followed by a singly modified form and a minor population consistent with a second ligand association.

ESI-MS analysis of the C313A and C313S mutants showed a mass increase consistent with Z34 modification under 1× treatment conditions in both variants (Figure S6A,B), indicating that ligand association is not dependent on Cys313.

Additional experiments performed at higher Z34 ratios (5×, 9×, and 12.5× equivalents per monomer) revealed further mass increases consistent with up to five ligand-associated species (Figure S6C), supporting the presence of additional modification sites at elevated ligand concentrations.

Together, these results identify Cys314 as the primary site of Z34 covalent modification, while indicating that additional cysteine residues can also be modified at higher ligand concentrations.

### Identification of labeled cysteine residues by MS after proteolytic digestion

To identify the location of the modified cysteine residues, ALDH1A3 was treated with Z34 and prepared for LC-MS/MS analysis. Following DMSO or Z34 treatment, the protein was denatured, reduced, alkylated, and digested with trypsin overnight and resulting peptides were analyzed by LC-Ms/MS. Data analysis revealed three Z34-modified cysteines: C52, C301, and the catalytic C314 (Fig. 4C-E). This result correlate with the intact mass spectrometry results showing multiple ligand adducts and also the labeling of the catalytic C314.

### Cryo-EM structure of the ALDH1A3-Z34 complex

To verify that ALDH1A3 was fully inhibited under the conditions used for structural studies, the enzyme was incubated overnight with increasing molar ratios of Z34 relative to the monomer (3×–12.5×) under conditions mimicking those used for cryo-EM sample preparation. Enzymatic activity measurements showed that all tested conditions resulted in nearly complete inhibition, with less than 0.45% residual activity (Figure S3B). A 3× inhibitor-to-monomer ratio was selected for structural studies to ensure active-site occupancy while minimizing nonspecific modification at higher ligand concentrations. Samples prepared under these conditions were subsequently used for cryo-EM analysis.

The cryo-EM map of the ALDH1A3-Z34 complex is of high quality, with an overall resolution of 2.26 Å, allowing the construction and refinement of a three-dimensional model with a map correlation of 84.6% (see Methods and Figure S7). In the map many solvent molecules are visible and most residues show well defined side chains. As reported previously for ALDH1A3 (32), the molecule is a tetramer with accurate D2 symmetry, each subunit having the characteristic fold of aldehyde dehydrogenases with continuous density spanning between residues 19 and 508.

There was some strong extra map close to the catalytic cysteine Cys314 (Figure 5), which was modeled as a Z34 molecule covalently bound, by its reactive acrylic carbon, to the cysteine sulfur atom. Interestingly this Z34 molecule presents two very different orientations with complementary occupancies (Figure 5A). The first orientation is towards the substrate pocket (57% occupancy in our refined model) (Figure 5B), while the second orientation is towards the NAD^+^ cofactor pocket (43% occupancy) (Figure 5C). These alternative orientations are achieved with essentially no rearrangements in the protein and a change of only 0.25 Å in the position of the sulfur atom from Cys314. The density in the substrate-binding pocket was not fully explained by the covalently bound Z34, and an additional non-covalently bound Z34 molecule was therefore modeled. This “non-covalently” bound Z34 has its acrylamide group away from Cys314 and making a hydrogen bond with the side chain of Trp189, while its two rings from the thiazole-thiophene moiety show about the same interactions as the covalently bound Z34 oriented towards the substrate pocket. Occupancy of this “non-covalently” bound Z34 is equal to the occupancy of the Z34 covalently bound and oriented towards the cofactor pocket. Therefore, ALDH1A3 subunits with Cys314 covalently bound to a Z34 in the cofactor pocket (43% in our refined model) contain also a “non-covalently” bound Z34 molecule in the substrate pocket. For the remaining ALDH1A3 subunits (57% in our model), the substrate pocket is occupied by the covalently bound Z34 oriented towards this pocket, while the cofactor pocket is empty (Figure 6). The cryo-EM map corresponds to the averaged of all subunits and consequently to the weighed averaged of the subunits displaying the two alternative orientations of the covalently bound Z34 molecule to Cys314. Biochemical results (see above) suggest that the percentages of each orientation could vary according to the relative concentrations of ALDH1A3 subunits and Z34 molecules and, when NAD^+^ is present, also to the competition for the binding in the cofactor pocket between Z34 and NAD^+^ molecules.

**Figure 5.**
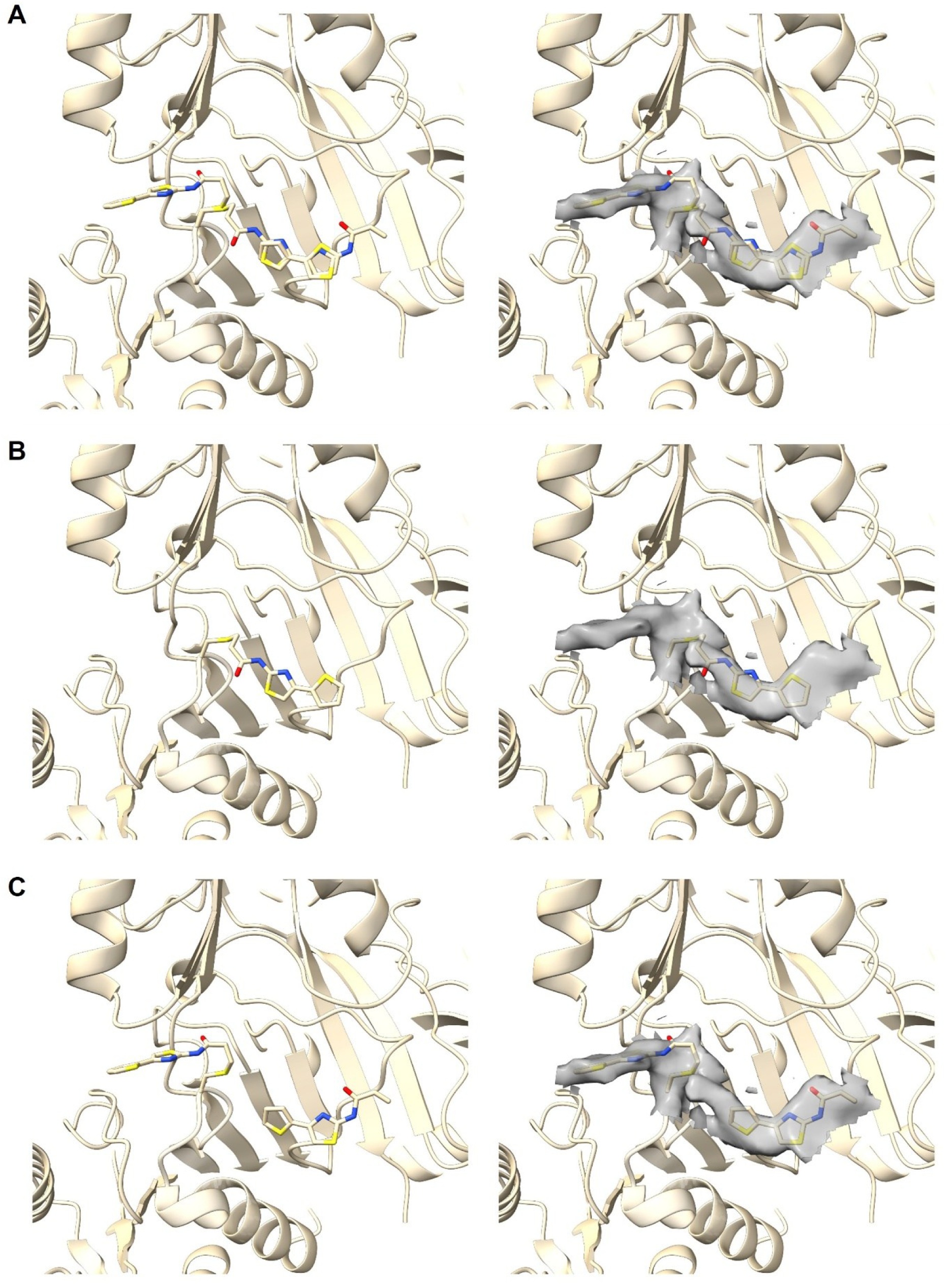
Structural representation of Z34 binding within the ALDH1A3 active site. The extra map surrounding Cys314 is shown as a semitransparent surface (right panels), whereas the corresponding atomic models are displayed in the left panels. The experimental map corresponds to the averaged mixture of two alternative covalent binding modes of Z34 to Cys314. **(A)** Final averaged model of the two alternative binding modes, each with approximately 50% occupancy. This combined model accounts for all the extra map observed around Cys314. **(B)** Representation of the first binding mode, in which a Z34 molecule is covalently bound to Cys314 while occupying the cofactor-binding pocket. A second Z34 molecule is non-covalently bound within the substrate-binding pocket, with its acrylamide warhead oriented away from Cys314. This structural state accounts for most of the experimental map but leaves additional unexplained map within the substrate-binding pocket. **(C)** Representation of the second binding mode, in which the covalently bound Z34 occupies the substrate-binding pocket. Likewise, this structural state accounts for only part of the experimental map, leaving additional map within the cofactor-binding pocket and part of the substrate-binding pocket.

**Figure 6.**
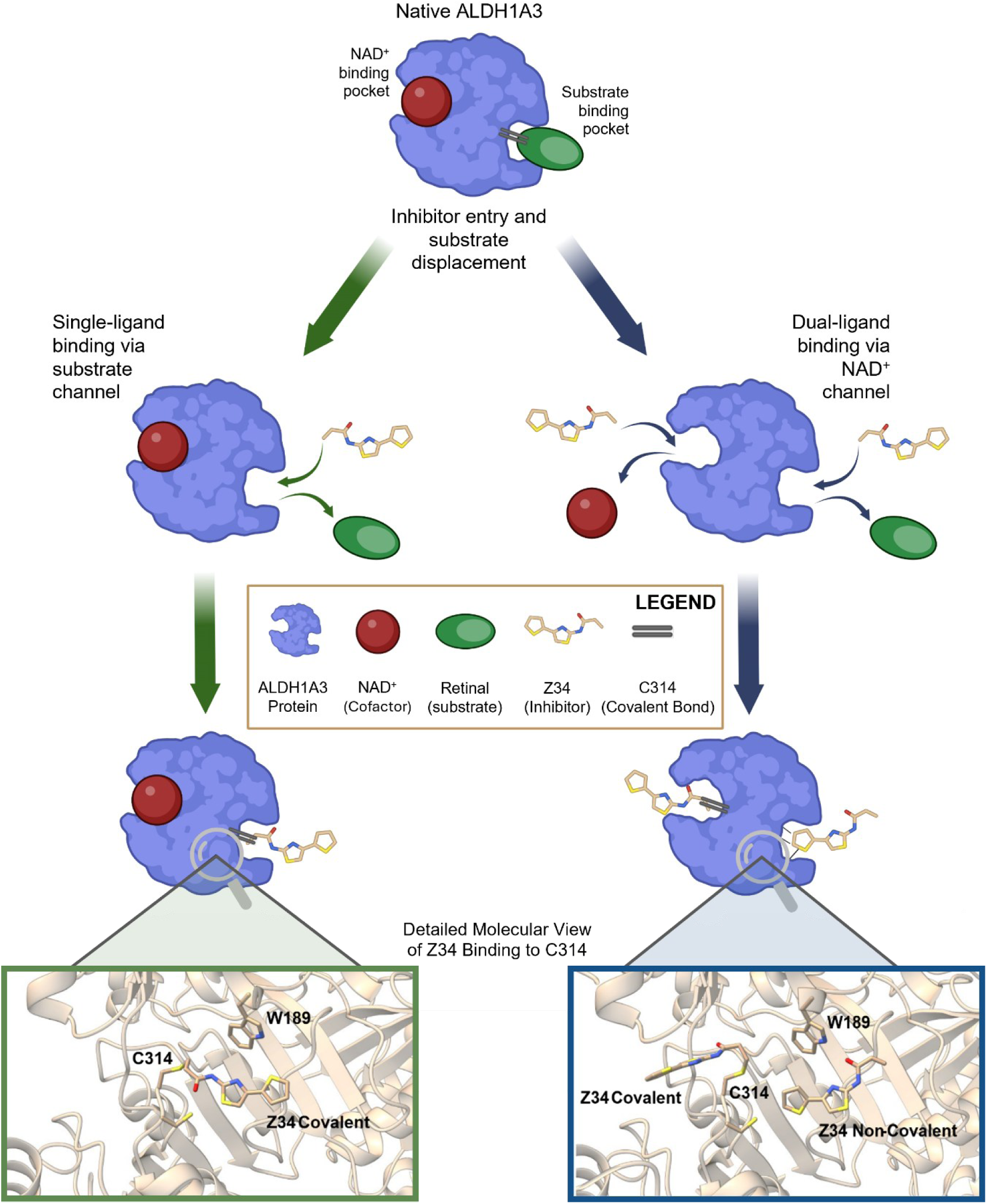
Binding alternatives of inhibitor Z34 to the catalytic residue Cys314 from ALDH1A3. Scheme to illustrate the two alternative covalent binding modes between Z34 and the catalytic Cys314. In the left, the covalently bound Z34 is oriented towards the substrate-binding pocket. In the right, the covalently bound Z34 is oriented towards the cofactor-binding pocket. In this second binding alternative another Z34 molecule is also found, although “non-covalently” bound, occupying the substrate pocket. This “non-covalently” bound Z34 has its warhead away from Cys314 making a hydrogen bond to Trp189. Created with BioRender.com

In two surface-exposed cysteines, Cys52 and Cys61, a small amount of extra map is visible, which has not been modelled and likely corresponding to the presence of some covalent complex with Z34 in these two residues. In turn, the better-defined extra map close to Cys301 has been modelled as a covalently bound Z34 molecule with full occupancy. In any case, these three cysteine complexes do not alter the structure of subunits and are away from the active center, which makes unlikely they could have a significant role in the inhibition of ALDH1A3 by Z34. Finally, it is worth mentioning that in the map there is not any hint of a covalent interaction between Cys313, the residue preceding the catalytic Cys314, and a Z34 molecule.

### Intrinsic tryptophan fluorescence reports on Z34 proximity to active-site residues

Given that our structural studies revealed that Z34 binds in close proximity to tryptophan residues located near the active site, we exploited intrinsic tryptophan fluorescence to probe local environmental changes upon ligand binding. Accordingly, intrinsic tryptophan fluorescence of ALDH1A3 (0.4 µM) was monitored upon incubation with 20 µM Z34 at 25°C (Figure 7B).

**Figure 7.**
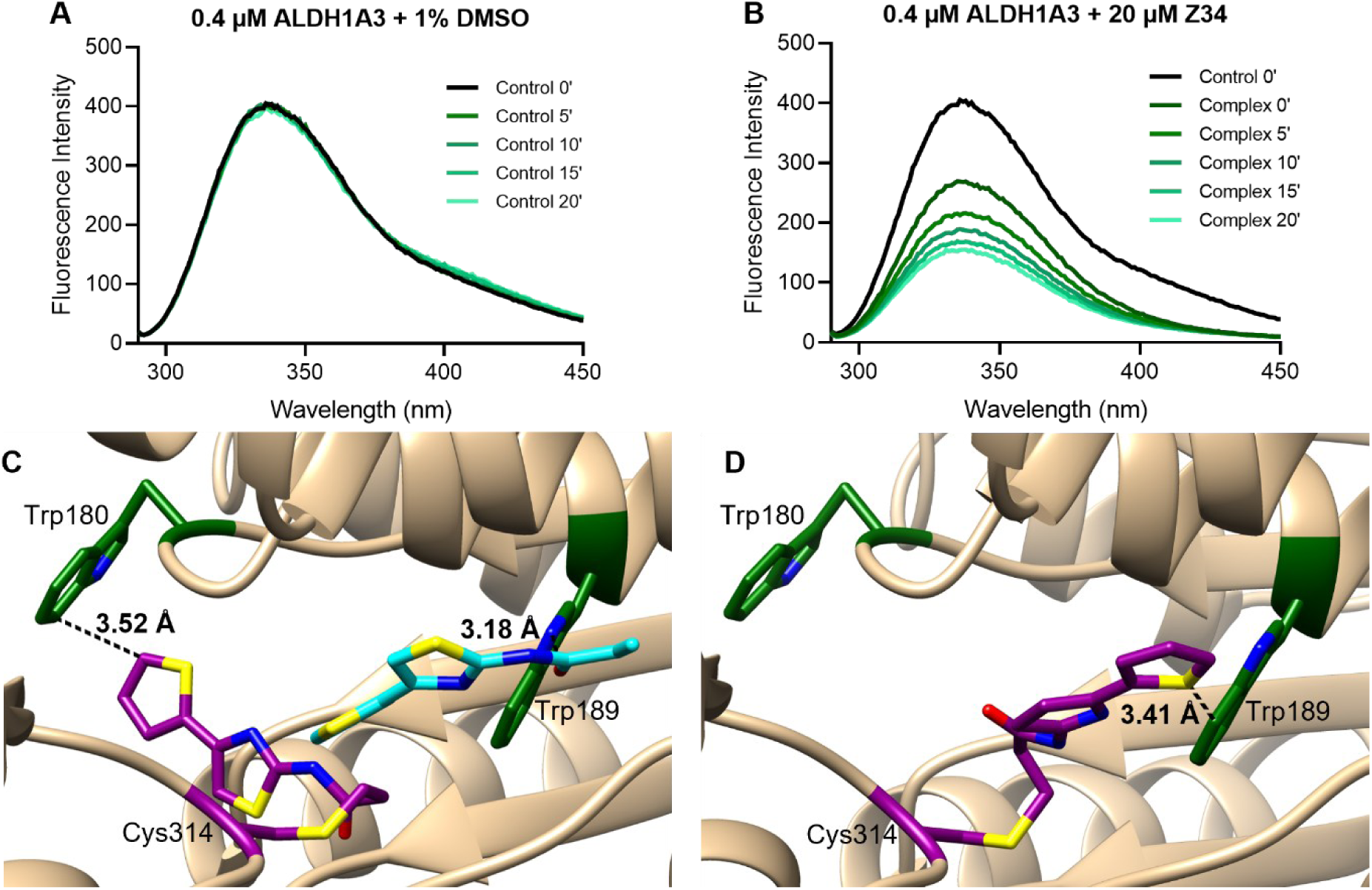
Structural basis of quenching of tryptophan fluorescence upon Z34 binding to ALDH1A3. **(A)** Maximum intrinsic tryptophan fluorescence emission of 0.4 µM ALDH1A3 in the presence of 1% DMSO (control) over a 0–20-min incubation period in the absence of NAD⁺. Protein samples were excited at 280 nm and emission maxima were monitored to assess potential changes in fluorescence intensity over time. **(B)** Maximum intrinsic tryptophan fluorescence emission of 0.4 µM ALDH1A3 in the presence of 20 µM Z34 over a 0–20-min incubation period, in the absence of NAD⁺. Protein samples were excited at 280 nm and emission maxima were recorded to monitor changes in fluorescence intensity upon inhibitor binding. **(C)** Structural representation of the ALDH1A3 active site showing Z34 binding via the NAD⁺ channel. Distances between nearby tryptophan residues (Trp180 and Trp189) and both covalently and non-covalently bound Z34 molecules are indicated, highlighting their close proximity (<4 Å). **(D)** Close-up view of Z34 bound via the substrate-binding channel, showing the proximity between the covalently bound inhibitor and Trp189. Distances between the indole ring and the ligand are indicated.

In the absence of inhibitor, the protein exhibited a stable maximum emission of 426.1 FU at 336 nm (Figure 7A). In contrast, addition of Z34 resulted in an immediate decrease in fluorescence intensity (0 min: 258.7 FU), which continued to decrease slightly over time (Table S2).

Given that the overall protein architecture remains largely unchanged in the cryo-EM structure of the ALDH1A3-Z34 complex, this decrease in fluorescence is likely explained by changes in the local environment of nearby tryptophan residues rather than large-scale conformational rearrangements. Structural analysis revealed that covalently bound Z34 molecules are located within 3.5 Å of Trp180 or 3.7 Å of Trp189 depending on the binding orientation, while a non-covalently bound Z34 molecule lies 3.4 Å from Trp189 (Figure 7C–D). The close proximity between the inhibitor and these residues provides a structural explanation for the rapid fluorescence quenching observed upon inhibitor addition.

### Effect of Z34 on cancer cells in culture

To evaluate the cellular activity of Z34, its effects were investigated in the human non-small cell lung cancer cell line A549 and the human glioblastoma cell line A172. Previous studies have reported expression of ALDH1A3 in both cell lines, together with additional ALDH family members (1,33,34).

Treatment with Z34 reduced cell viability in both models (Figure 8A,C). In A549 cells, EC₅₀ values were 16.6 ± 4.3, 16.4 ± 3.9, and 20.3 ± 1.3 µM after 24, 48, and 72 h, respectively (Figure S8A and Figure 8A). In A172 cells, EC₅₀ values were 17.7 ± 3.2, 16.6 ± 2.8, and 12.3 ± 0.8 µM at the corresponding time points (Figure S8B and Figure 8C).

**Figure 8.**
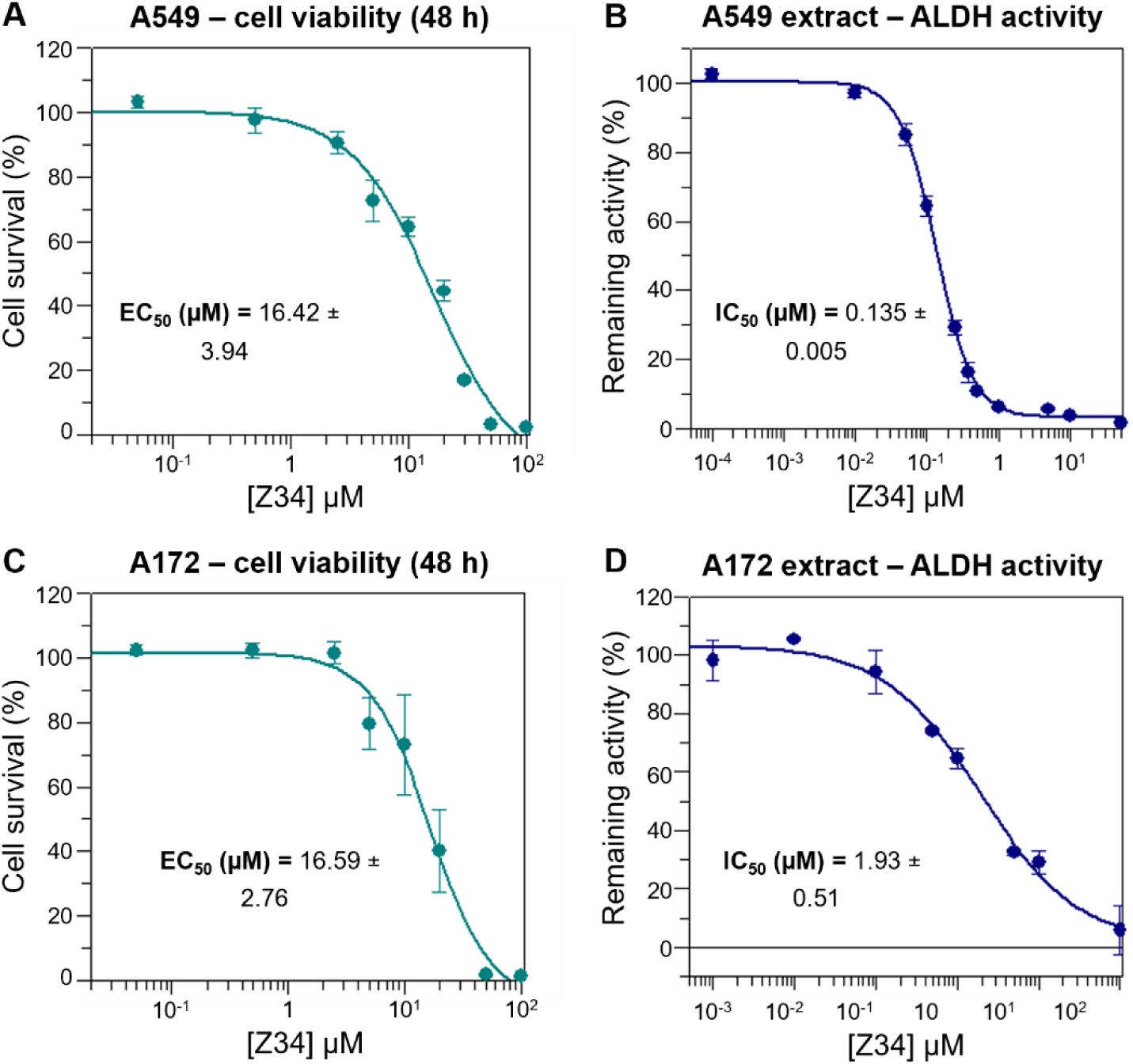
Cellular activity of Z34 in cancer cell models expressing ALDH isoforms. **(A)** Effect of Z34 on A549 cell viability after 48 h treatment. Dose–response curve was used to determine EC₅₀ value. Data represents SD from three independent experiments; each performed with three technical replicates per condition. **(B)** Inhibition of ALDH activity by Z34 in A549 cell extract under conditions favoring ALDH1A3 activity (50 mM HEPES, 50 mM MgCl₂, pH 8.0, hexanal 250 µM). Extract was pre-incubated with Z34 for 20 min at 25°C prior to activity measurement. **(C)** Effect of Z34 on A172 cell viability after 48-h treatment. Dose–response curve was used to determine EC₅₀ value. Data represents SD from three independent experiments; each performed with three technical replicates per condition. **(D)** Inhibition of ALDH activity by Z34 in A172 cell extract under ALDH1A3-like conditions, as described in panel B.

To determine whether Z34 inhibits ALDH activity in a cellular context, enzymatic activity was measured in cell extracts following a 20 min pre-incubation with inhibitor. In A549 extracts, Z34 inhibited ALDH activity with an IC₅₀ value of 0.135 ± 0.005 µM under conditions optimized for ALDH1A3 activity (Figure 8B). In A172 extracts, inhibition was also observed, yielding an IC₅₀ value of 1.93 ± 0.51 µM (Figure 8D). Notably, the potency observed in cellular extracts was comparable to or greater than that measured using recombinant ALDH1A enzymes, with A549 extracts displaying particularly high sensitivity to Z34.

Together, these findings extend the biochemical characterization of Z34 to a cellular context, demonstrating potent inhibition of endogenous ALDH activity in cellular extracts and an associated reduction in cell viability in two cancer cell models.

## DISCUSSION

Our findings demonstrate that Z34 is a covalent irreversible inhibitor targeting the catalytic Cys314 of ALDH1A3. Covalent complex formation increased the thermal stability of both the binary ALDH1A3-Z34 and ternary ALDH1A3-NAD⁺-Z34 complexes, supporting the formation of stable inhibitor-bound states.

Z34 contains a classical acrylamide (prop-2-enamide) warhead attached to a thiazole-thiophene scaffold. Acrylamide electrophiles are widely used in targeted covalent inhibitor design because they react with nucleophilic cysteine residues through 1,4-Michael addition (35,36). Recently, the KRAS inhibitor ARS-1620 was also identified as an off-target covalent inhibitor of ALDH1A3 through modification of Cys314 (37), and docking studies supported a Michael addition mechanism. In contrast, the present cryo-EM structure directly visualizes the covalent thioether linkage formed between Z34 and Cys314 and further reveals two mutually exclusive binding conformations within the ALDH1A3 active site, providing experimental structural insight into covalent ligand recognition by this enzyme.

The three-dimensional structure of the ALDH1A3-Z34 complex, determined by Cryo-EM at 2.26 Å resolution, provides the structural basis for the biochemical observations described above. To our knowledge, this first Cryo-EM structure of human ALDH1A3 reveals two mutually exclusive binding conformations within the active site: one Z34 molecule occupies the substrate-binding pocket while the other locates oppositely in the cofactor-binding site. The Z34 molecule bound in the NAD^+^-binding site is compatible with the simultaneous presence in the substrate-binding pocket of a non-covalently bound Z34 molecule, oriented with its acrylamide group facing the substrate entry channel. To our knowledge, this represents the first high-resolution structure directly visualizing acrylamide-mediated covalent modification of the catalytic cysteine in an ALDH1A enzyme. The thiazole-thiophene scaffold establishes extensive hydrophobic interactions, particularly with conserved tryptophan residues, providing a structural explanation for the high affinity of Z34 for the catalytic pocket.

These alternative binding modes (to the substrate- or cofactor-binding site) are unprecedented within the ALDH family. Although the experimental conditions favored Z34 binding because inhibitor screening and pre-incubation were performed in the absence of NAD⁺, covalent ligand binding extending into the cofactor-binding site has only rarely been described. Covalent linkage between the catalytic cysteine and the nicotinamide ring has previously been predicted for ALDH2 (38) and observed in ALDH1L1 and betaine aldehyde dehydrogenase (39,40), while brominated cofactor analogues have also been shown to label the catalytic cysteine (41). Furthermore, several studies suggest that, when the substrate entrance is sterically blocked, ligands may access the active site through the cofactor channel (42–45). The present structure extends these observations by directly visualizing two alternative covalent binding trajectories for a small-molecule inhibitor within a human ALDH enzyme.

Notably, despite its immediate proximity to the catalytic residue, Cys313 was not detectably modified by Z34. This conclusion is supported by mutagenesis, mass spectrometry and cryo-EM, all of which consistently identify Cys314 as the primary site of covalent modification. These observations indicate that productive modification depends on precise molecular recognition within the active site rather than on simple accessibility of a cysteine thiol. In contrast, Cys301 was identified as a secondary site of modification by both cryo-EM and MS/MS analyses, consistent with the additional covalent adducts detected at elevated inhibitor concentrations. Interestingly, modification of Cys301 was also reported in the recent proteomic analysis of ARS-1620-treated ALDH1A3 (37), suggesting that this residue may represent a recurrent secondary target for cysteine-reactive electrophiles. Importantly, despite containing a reactive acrylamide warhead, Z34 did not inhibit three representative cysteine proteases included in the primary profiling panel, further supporting the notion that productive covalent inhibition requires both chemical reactivity and favorable molecular recognition. Collectively, these findings indicate that covalent modification of Cys314 represents the primary inhibitory mechanism, whereas modification of other cysteine residues corresponds to secondary labeling events.

Comparison with previously reported irreversible ALDH inhibitors highlights several distinctive features of Z34 (Table 2). Based on its *k_inact_/K_i_* value, Z34 ranks among the most efficient covalent inhibitors reported for the ALDH1A family despite its relatively simple chemical scaffold. Covalent ALDH inhibitors reported to date comprise diverse electrophilic warheads and reaction mechanisms, including disulfide formation, thioacylation and Michael addition, and differ substantially in isoform selectivity. Z34 and EN40 represent the first ALDH inhibitors for which an acrylamide warhead has been deliberately exploited to generate a stable thioether linkage with the catalytic cysteine, although EN40 selectively targets ALDH3A1 (46). In addition, the recent identification of the KRAS inhibitor ARS-1620 as an off-target covalent modifier of ALDH1A3 further supports the suitability of acrylamide electrophiles for targeting this enzyme family (37). Together, these observations establish acrylamide warheads as an effective strategy for irreversible inhibition of therapeutically relevant ALDH isoforms.

**Table 2.**
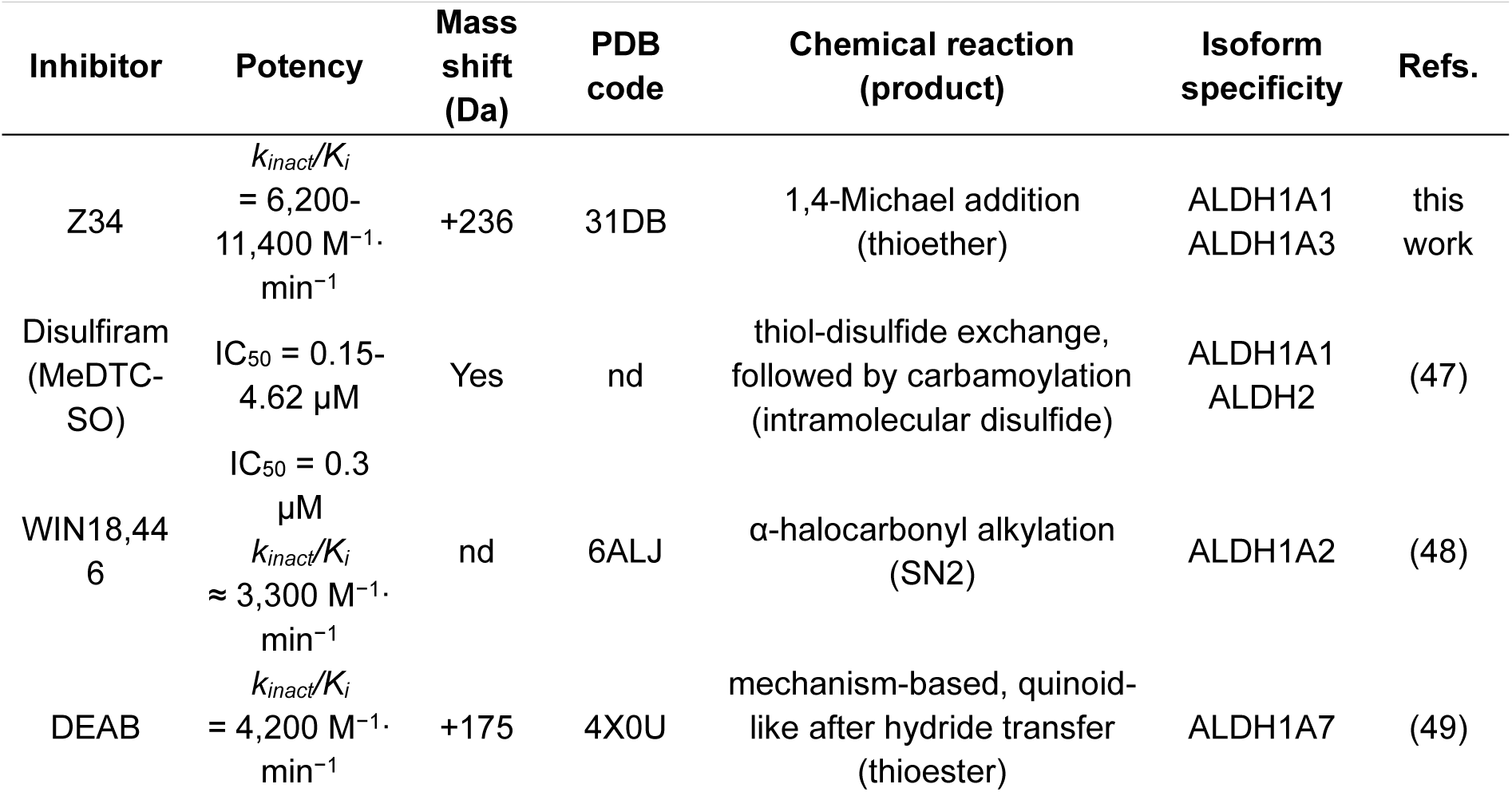
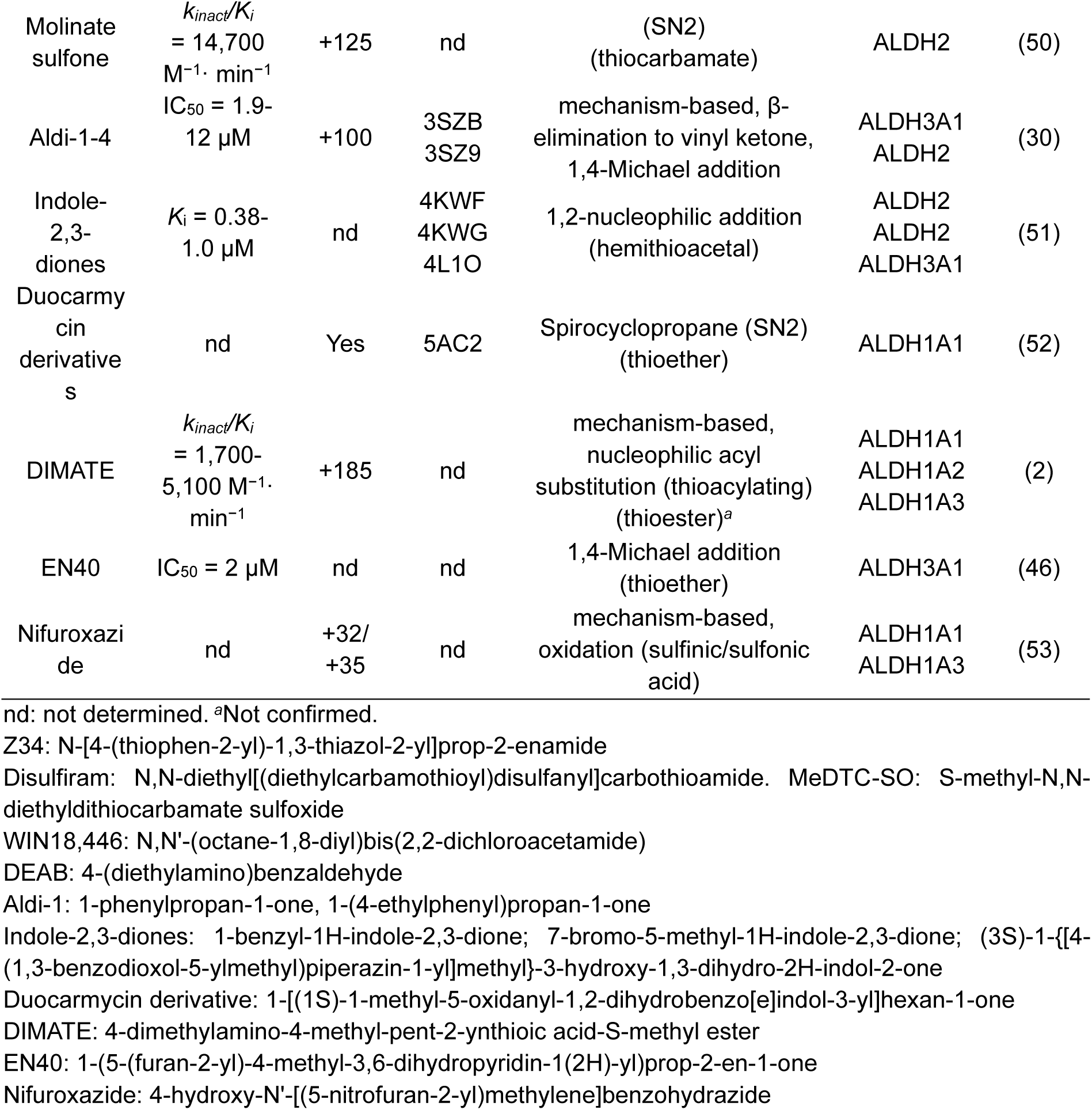
Properties of compound Z34 compared to those of other covalent ALDH inhibitors binding to the active-site cysteine residue.

An additional contribution of this work is methodological. High-throughput screening of ALDH inhibitors can be technically challenging when based on the physiological dehydrogenase reaction, as it requires monitoring NADH production using aldehyde substrates that are often volatile or poorly suited for high-throughput assay formats. The fluorogenic esterase assay developed here, based on a series of 4-methylumbelliferyl esters identified for the first time as ALDH substrates, provided a robust platform for primary screening. Subsequent mechanistic characterization using the physiological aldehyde dehydrogenase reaction confirmed that the identified hits also inhibited the native catalytic activity. Collectively, these findings establish ALDH esterase activity as a practical surrogate for ALDH inhibitor discovery and expand the experimental toolbox available for investigating ALDH enzymes through a simple fluorogenic platform for future mechanistic studies and inhibitor development.

Although the primary focus of this work was mechanistic characterization at the recombinant enzyme level, inhibition of endogenous ALDH activity in cancer cell extracts together with the associated reduction in cell viability demonstrates that the biochemical properties of Z34 are retained in a cellular environment. These observations provide an initial proof of cellular activity while supporting the relevance of the mechanistic findings obtained *in vitro*.

Covalent enzyme inhibition has re-emerged as a powerful strategy in chemical biology and drug discovery because durable target engagement can produce sustained pharmacological effects beyond compound exposure (17,18). Within this context, the combined biochemical, mass spectrometric, structural and cellular analyses presented here provide a molecular framework for understanding Z34 recognition by ALDH1A enzymes, define the structural determinants governing selective modification of the catalytic cysteine, and establish a foundation for the rational design of next-generation covalent inhibitors targeting this therapeutically relevant enzyme family.

## CONCLUSIONS

This work identifies the Enamine compound Z3405279217 (Z34) as a novel covalent irreversible inhibitor of the human ALDH1A3 and ALDH1A1 isoforms through an integrated workflow combining a newly established 4-methylumbelliferyl esterase-based high-throughput screening assay with comprehensive biochemical, structural, and cellular characterization. Following hit identification, mechanistic studies based on the physiologically relevant aldehyde dehydrogenase activity demonstrated that Z34 is a time-dependent inhibitor whose initial interaction with ALDH1A3 is modulated by NAD⁺ and reducing agents, while retaining high inhibitory potency after covalent inactivation. Thermal shift assays, mass spectrometry, and high-resolution cryo-EM identified the catalytic Cys314 as the primary site of covalent modification and revealed that Z34 adopts two mutually exclusive binding conformations within the active site, providing a structural explanation for its inhibitory mechanism. Although additional cysteine residues became modified at elevated inhibitor concentrations, these events represent secondary interactions rather than the primary mechanism of enzyme inhibition. Finally, Z34 effectively inhibited endogenous ALDH activity in cancer cell extracts and reduced cell viability in two cancer cell models, demonstrating that the compound remains functionally active in a cellular environment. Together, these findings establish Z34 as a valuable chemical probe for investigating ALDH1A enzymes and provide the first structural framework for understanding and rationally designing covalent inhibitors targeting the ALDH1A family.

## MATERIALS AND METHODS

### ALDH purification and lyophilization

Recombinant human ALDH1A1 and ALDH1A3 (wild-type), as well as ALDH1A3 active-site mutants (C314S, C313A, and C313S), containing an N-terminal (His)_6_ tag, were overexpressed and purified as previously described (54,55). Purification was performed by Ni²⁺-charged agarose affinity chromatography using a HisTrap HP 5 mL column (Cytiva) on an ÄKTA™ fast protein liquid chromatography system. Recombinant proteins were stored at – 20°C in 20 mM Tris-HCl, 250 mM NaCl, 5 mM DTT, pH 8.0.

For lyophilization, ALDH1A3 was supplemented with trehalose as a cryoprotectant at a 10-fold mass excess relative to protein (10:1, w/w) prior to freezing in liquid nitrogen. Lyophilization was performed for 24 h at 25 mTorr and –105°C after freezing in liquid nitrogen using a VirTis benchtop lyophilizer equipped with a Welch 8917 vacuum pump.

### Sample preparation and storage

Lyophilized ALDH1A3 was stored at −20°C upon receipt. For assay sample preparation, ALDH1A3 was dissolved in MilliQ water to 36 µM (determined by BCA assay) and the protein stock solution was aliquoted, flash-frozen, and stored at −80°C. To avoid multiple freeze-thaw cycles, protein stock solutions thawed from −80°C were stored at 4°C for further use.

### ALDH Esterase Assay

Lyophilized ALDH1A3 was resuspended in Milli-Q water to prepare a 36 μM stock solution and stored as 10 μL aliquots at −80°C to minimize freeze–thaw cycles. Esterase activity was measured by diluting freshly-thawed ALDH1A3 stock solution into assay buffer containing 50 mM HEPES (pH 8.0), 50 mM MgCl₂, and 10⁻⁶% (v/v) Triton X-100 to achieve a final enzyme concentration of 0.3125 μM. Five different 4-methylumbelliferyl (4-MU)-based fluorogenic substrates with varied carbon lengths (2c, 4c, 7c, 8c, and 10c) were used for substrate activity. The substrate stocks were prepared by diluting 10 mM DMSO stocks (56) in assay buffer to achieve a final substrate concentration of 250 μM. 8 µL Enzyme stock solution was mixed with 2 μL substrate stock solution in a Greiner 384-well low-volume, flat-bottom, all-black plate (Greiner 78407625). Fluorescence measurements (λ_ex_ = 335 nm, λ_em_ = 450 nm) were recorded at least every 3 min for a total of 1 hour using a Cytation 3 Multi-Mode Reader (BioTek, Winooski, VT). Turnover rates were determined during the linear phase of the reaction (40 min) prior to substrate depletion and expressed as RFU/min in GraphPad Prism. The reaction rates for each substrate were normalized by subtracting background hydrolysis rates measured in reaction buffer in the absence of protein.

### High-throughput screening

ALDH1A3 was screened against the commercial Enamine Cysteine-focused covalent library containing 3,200 compounds available at the High-Throughput Screening Knowledge Center (HTSKC) at Stanford University. A Beckman Echo 655 acoustic liquid handler was used to transfer 50 nL from source plates containing 20 mM DMSO stocks into assay plates to achieve a final concentration of 100 μM for the screening. All screening assays were conducted using 250 nM ALDH1A3 and 50 μM methylumbelliferyl caprylate (4-MUC) as the final substrate concentrations.

For initial screening, 0.250 μM ALDH1A3 enzyme was incubated with 100 μM of each compound for 1 hr. N-ethylmaleimide (NEM) was used as the positive inhibition control, whereas DMSO served as the negative control. Following incubation 50 μM substrate was added to each well, and fluorescence was monitored using the plate reader in endpoint mode at λ_ex_ = 365 nm and λ_em_ = 455 nm for 1 h. Assay quality was evaluated using the Z′ factor according to Equation 1 (57).

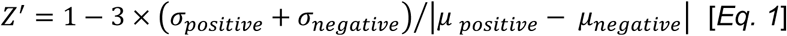

Where *σ* refers to the standard deviation of fluorescence readout from positive or negative groups, and *μ* refers to the standard deviation of fluorescence readout from positive or negative groups. The percentage of inhibition was calculated using Equation 2.

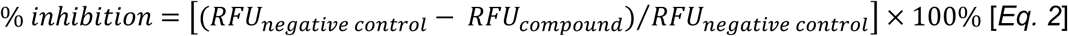

Where *RFU_compound_* fluorescence values from wells containing test compounds represent fluorescence values from wells without inhibitor, and *RFU_compound_* refers to the fluorescence reading of the enzymatic reaction in the presence of the compound.

Compounds with more than 95% inhibition were selected from the library for dose-response tests to calculate IC_50_ values. The dose-response setup is the same as in the initial screening assay, except for different inhibitor concentrations.

### Compounds from Enamine library

The Enamine cysteine-focused covalent library (3,200 compounds) comprises diverse electrophilic scaffolds designed for covalent engagement of protein cysteine residues. The collection includes acrylamides, dimethylamine-substituted acrylamides, chloroacetamides, 2- chloropropionamides, chlorofluoroacetamides, and butynamides, selected to balance reactivity and selectivity for hit identification in covalent screening campaigns. It was built by combining well-validated electrophilic warheads with structure-based and using the rule-of-three criteria (molecular weight less than 300 Da, no more than 3 hydrogen bond donors, no more than 3 hydrogen bond acceptors, and cLogP of 3 or less) to minimize overly reactive or promiscuous compounds.

### Z34 Compound: Z3405279217

N-[4-(thiophen-2-yl)-1,3-thiazol-2-yl]prop-2-enamide (CAS number 2305402-77-9) was obtained from Enamine US Inc (Monmouth Jct., NJ 08852,USA).

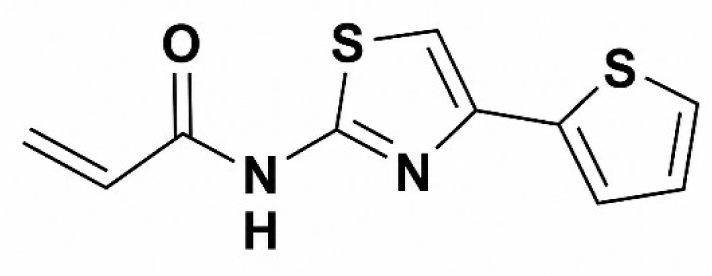

### Dehydrogenase activity assay

Enzymatic activity was monitored using a Cary Eclipse Varian fluorometer by measuring NADH fluorescence at 460 nm (excitation at 340 nm) with 10 nm slit bandwidths. Reactions were performed in 50 mM HEPES buffer pH 8.0 containing 50 mM MgCl₂ for ALDH1A3, or 50 mM HEPES buffer pH 8.0 containing 0.5 mM EDTA for ALDH1A1, at 25°C maintained by an external thermostatic water circulator.

Assays contained 500 μM NAD⁺ and were initiated by addition of hexanal at varying concentrations. Fluorescence was monitored for 600 s. Conversion of fluorescence units into absolute reaction rates using 5 μM NADH as an internal standard was performed as previously described (2). Kinetic parameters for both enzymes were determined from measurements over a range of hexanal concentrations.

### Inhibition kinetics assays

Initial inhibition screening was performed using the dehydrogenase activity assay described above to evaluate the inhibitory activity of Z34 at 10 μM against ALDH1A1 and ALDH1A3. Z34 was dissolved in DMSO and assayed at a final concentration of 1% (v/v). Enzyme-inhibitor complexes were pre-incubated in the absence of NAD⁺, and reactions were initiated by addition of 500 µM NAD⁺ and substrate at concentrations corresponding to both saturating and *K_m_* values. Reactions were incubated for 5 and 20 min to assess potential time-dependent effects. Screening experiments were performed in duplicate, and values were reported as the mean ± standard deviation using GraphPad Prism 8.0 (GraphPad Software, Boston, MA, USA) as previously described (58).

IC_50_ values were determined at saturating substrate concentrations after 20 min incubation, by non-linear regression using GraFit 5.0 (Erithacus Software, UK) as described previously (58). Experimental values were represented as the mean ± standard deviation from two independent measurements, while IC_50_ value was expressed as the mean ± standard error.

### Determination of *k_obs_* and *k_inact_/K_i_*

To determine the time-dependent inhibitory potency of Z34 against ALDH1A1 and ALDH1A3, different inhibitor concentrations were incubated with the enzymes for 0, 1, 2, 5, 10, 20, 30, 45, and 60 min at 25°C in the appropriate reaction buffer prior to the addition of 500 μM NAD⁺, the NADH internal standard, and saturating substrate concentrations. Inhibitor solutions were assayed at a final DMSO concentration of 1% (v/v). Enzyme concentrations were maintained 50- to 500-fold lower than that of the substrate in all assays.

Apparent rate constants (*k_obs_*) were obtained at saturating substrate concentrations by fitting the data using GraFit 5.0, as previously described (2). The dependence of *k_obs_* on inhibitor concentration was used to calculate *k_inact_/K_i_*. Data are presented as the mean ± standard deviation from duplicate measurements, and *k_obs_* and *k_inact_/K_i_* are reported as mean ± standard error.

### Competitive effect of NAD^+^ on ALDH1A3-Z34 interaction

Competition experiments were performed to evaluate the effect of NAD⁺ on Z34-mediated inhibition of ALDH1A3. Enzyme–inhibitor complexes were pre-incubated with Z34 (75 µM and 10 µM) in the presence of increasing NAD⁺ concentrations (0, 1, 20, 100, 500, and 1000 µM) for 20 min. Reactions were carried out in 50 mM HEPES buffer pH 8.0 containing 50 mM MgCl₂ at 25°C.

After the pre-incubation period, reactions were adjusted to a final NAD⁺ concentration of 500 µM (except for samples already containing 500 or 1000 µM NAD⁺). Assays were supplemented with 5 µM NADH as an internal standard and initiated by addition of hexanal. For wild-type ALDH1A3, enzymatic activity was measured either at saturating substrate concentration (250 µM hexanal) or under low-substrate conditions well below the *K_m_* (2.5 µM hexanal; *K_m_* ≈ 15 µM). For the C313A mutant, activity measurements were performed only under saturating substrate conditions (60 µM hexanal).

Enzymatic activity was monitored as described above. Experiments were performed in duplicate and data were analyzed using one-way ANOVA with multiple comparisons against the 0 µM NAD⁺ control using GraphPad Prism 8.0 software. Values are reported as mean ± standard deviation.

### Thermal stability assay

The thermal stability of ALDH1A3 in the presence of Z34 was assessed using a fluorescence-based thermal shift assay with SYPRO Orange (Invitrogen), following the method described by Niesen et al. (59). Protein (1 µM) was diluted in 10 mM HEPES buffer, pH 7.5, containing 150 mM NaCl. A 4× stock of TSA buffer containing SYPRO Orange was freshly prepared and added to each well, yielding a final 1× buffer concentration and a dye concentration of 1:1000 in the assay. Reactions were performed in the absence or presence of 500 µM NAD⁺, with Z34 or DMSO added to achieve final concentrations ranging from 1 to 100 µM (1% v/v DMSO).

Reactions were performed in a final volume of 10 µL per well in 96-well PCR plates (Bio-Rad). Fluorescence was recorded on a CFX96 Touch Real-Time PCR System (Bio-Rad) during a thermal ramp from 25°C to 75°C, increasing 0.3°C every 60 s.

Melting temperature (Tm) was determined by fitting the fluorescence data to a Boltzmann sigmoidal equation using GraphPad Prism 8.0. Each treatment at each inhibitor concentration was performed in triplicate, and the experiment was independently repeated. Tm values were presented as the mean ± standard deviation, and most curve fits yielded R² values above 0.8.

### ESI-TOF mass spectroscopy

Wild-type ALDH1A3 and C313S, C313A, and C314S mutants were diluted to 8.4 µM in 100 mM ammonium bicarbonate (pH 7.5). Protein concentrations were calculated based on the molecular weight of the tetrameric form of each variant. Z34 stock solutions (100×) were prepared in DMSO to give a final DMSO concentration of 1% (v/v). Fresh protein samples were incubated with Z34 at protein:ligand molar ratios of 1:4, 1:8, and 1:12, corresponding to 1, 2, and 3 equivalents of inhibitor per monomer, respectively, for 30 min at 25°C. Control samples contained DMSO only. For the C313S and C313A mutants, only control and 1:4 treatments were analyzed.

Samples were analyzed by direct injection without column under near-neutral conditions (pH 7.0) using a running buffer consisting of acetonitrile/water 0.1% formic acid (80:20, v/v) mobile phase. For each analysis, 10 µL of a 1:2 diluted protein solution was injected at a flow rate of 100 µL · min⁻¹.

Electrospray ionization positive mass spectrometry (ESI-TOF-HRMS) was performed on a MicroTOF-Q following instrument: a UHPLC Elute+ coupled to a timsTOF Pro2 system (Bruker Daltonics) and controlled using Bruker Compass HyStar 6.3 software. Instrument parameters were set as follows: capillary counter-electrode voltage 3500 V, m/z range 500–3000, dry temperature 180°C, and dry gas flow 6 L · min⁻¹. Raw spectra were processed and deconvoluted using the Maximum Entropy algorithm implemented in Bruker Compass DataAnalysis software (Bruker Daltonics). Reported molecular masses correspond to the monomeric species obtained after deconvolution.

### Identification of labeled cysteine residues by MS/MS after proteolytic digestion

ALDH1A3 (36 µM in 20 mM Tris, 500 mM NaCl, 5 mM DTT, pH 8) was diluted to 2 µM with PBS (14 µL total) and 1 µL of DMSO, or Z34 (150 µM, final 10 µM) was added and incubated for 30 min at room temperature. Samples were diluted to 6 M in 50 mM NH_4_HCO_3_ pH 8, reduced with 10 mM TCEP (fresh 100 mM stock in H2O) for 20 min, alkylated with 25 mM IAA (fresh 250 mM stock in H_2_O). Samples were diluted to 2 M urea with 50 mM NH_4_HCO_3_ pH 8 and digested for 5 h with tryspin (1 µL of 0.5 mg/mL) at 37°C in the presence of the 1 mM CaCl_2_. Samples were acidified with 1 vol. of isopropanol 1% TFA and desalted using styrenedivinylbenzene reverse-phase sulfonate (SDB-RPS) StageTips as described previously (60). Briefly, samples were loaded on a 200 µL StageTip containing two SDB-RPS disks and centrifuged at 1500 × g for 8 min. This was repeated until all the sample was loaded on the StageTip. StageTips were washed three times with 200 µL of isopropanol 1% TFA at 1500 × g for 8 min, then eluted with 100 µL of 80% MeCN, 19% water, and 1% ammonia and dried. Peptides were resuspended in water with 0.1 % formic acid (FA) and analyzed using nanoElute 2 coupled to a TimsTOF HT mass spectrometer (Bruker Daltonics). The chromatography column consisted of a 25 cm long, 150 μm i.d. microcapillary packed with 1.5 μm C18 particles (Bruker Daltonics/Pepsep) and was capped by a 20 μm emitter (Bruker Daltonics). LC solvents were 0.1% FA in H_2_O (Buffer A) and 0.1% FA in MeCN (Buffer B). Peptides were eluted into the MS at a flow rate of 500 nL/min. over a 30 min. linear-gradient (5-35 % Buffer B) at 50 °C. Data was acquired via dda-PASEF using 10 PASEF ramps per one MS1 scan and fragmented ions with charges 0 to 5. The method covered m/z 100-1700 and scan range between 0.6 to 1.6 1/K0. Ramp time was set to 100 ms and total cycle lasted 1.17 s. Collision energy was set as a linear increase from 20 eV at 1/K0 = 0.6 to 59 eV at 1/K0 = 1.6. The MS data was analyzed with Fragpipe (V22) (61,62) and searched against the Human proteome and common list of contaminants (generated in Fragpipe). Peptide and fragment mass tolerance was set to 20 ppm. The minimum peptide length was set to 6 amino acids with a mass range of 500 to 5000 Da. Digestion was set to semi. Oxidation of methionine, N- terminal acetylation, carbamidomethylation of cysteine and Z34 (+236.0078 Da) on cysteine were set variable modifications. PSM validation was performed with Peptide Prophet with the --decoyprobs --ppm --accmass --nonparam –expectscore parameters. Quantification was performed using IonQuant.

### Cryo-EM structure of the ALDH1A3-Z34 complex

Purified ALDH1A3 (16.78 µM tetramer; 67.14 µM monomer equivalent) was incubated overnight at 25°C with 201 µM Z34, corresponding to three equivalents of inhibitor per monomer. The sample was subsequently centrifuged at 20,000 × g for 30 min at 4°C, filtered through a 0.45 µm membrane, and purified by size-exclusion chromatography using a HiLoad 16/600 Superdex 200 pg column (Cytiva) connected to an ÄKTA fast protein liquid chromatography system. Fractions corresponding to the ALDH1A3-Z34 complex were pooled and diluted to 0.6 mg/mL in 20 mM Tris-HCl, 250 mM NaCl, 0.01% Brij-35, pH 8.0 for cryo-EM analysis.

For single-particle cryo-electron microscopy (cryoEM), 3 µL drop of purified ALDH1A3-Z23 complex was applied to glow-discharged C-Flat™ holey carbon grids (CF-1.2/1.3; 300 mesh), blotted for 4 s using a blot force of −3 and vitrified in liquid ethane using a Vitrobot Mark IV (Thermo Fisher Scientific) set to 4°C and 95% humidity. Cryo-EM grids were stored in liquid nitrogen until data collection.

Movies of ALDH1A3-Z34 complex were collected on a Titan Krios electron microscope operated at 300 kV (Diamond-eBIC). Imaging was performed using EPU at a nominal magnification of ×130,000 (0.65 Å/pixel) at 300 kV Titan Krios (Thermo Fisher Scientific) electron microscope equipped with K3 direct electron detector (Gatan). A total of 30,345 movies were collected. The camera was operated in counting mode with a dose rate of 0.579 electrons per Å^2^ · s^−1^, resulting in a total dose of 44.6 electrons per Å^2^. Defocus values ranged from −0.7 to −1.8 µm (Table S1).

CryoSPARC v4a was used to process the cryoEM data. Particles were selected with the Template picker using a particle diameter of 100 Å. Particles were extracted and classified in 2D. For the final processing, the 2D particles containing the complex were selected, remaining a total of 2,919,368 particles which were used to generate an ab initio reconstruction with three classes followed by a subsequent non-uniform heterogeneous refinement with the largest class. Finally, a model with a resolution of 2.4 Å was obtained. A second processing stage was carried out by taking the particles used in the non-uniform refinement and subjecting them to reference-based motion correction, during which a total of 8,432 particles were excluded. The accepted particles were then reprocessed through non-uniform refinement, resulting in a final map at 2.26 Å (Figure S7B).

Docking, tracing and refinement of all the structures was performed alternating interactive and automatic cycles with programs Coot (63) and Phenix (64). The final refined structure has been deposited in the PDB with code: 31DB and the map in the EMDB with code: 58311 (Figure S7F).

### Intrinsic tryptophan fluorescence

Intrinsic tryptophan fluorescence was monitored using a Cary Eclipse Varian fluorometer. Measurements were performed in a 1 mL quartz cuvette at 25°C maintained by an external thermostatic water circulator. The excitation wavelength was set to 285 nm and emission spectra were recorded between 300 and 450 nm using excitation and emission slit bandwidths of 10 and 5 nm, respectively.

To monitor ALDH1A3-Z34 interactions, ALDH1A3 (0.4 µM) was incubated with Z34 (20 µM) in 50 mM HEPES buffer pH 8.0 containing 50 mM MgCl₂. Emission spectra were collected at 0, 5, 10, 15, and 20 min after addition of the inhibitor. Control measurements were performed in the presence of 1% (v/v) DMSO under identical conditions.

### Cell culture and viability assays in cancer cell lines

The human non-small cell lung cancer cell line A549 and the human glioblastoma cell line A172 (ATCC) were cultured in high-glucose DMEM (Dominique Dutscher) supplemented with 10% fetal bovine serum (FBS; Fisher Scientific) at 37°C in a humidified atmosphere containing 5% CO₂.

Cell viability was assessed using a PrestoBlue™-based assay essentially as previously described (1). Cells were seeded in 96-well plates (Falcon, Corning) at a density of 3,000 cells per well and allowed to adhere overnight. Subsequently, cells were treated with increasing concentrations of Z34 (0.1–100 µM) for 24, 48, or 72 h. The final DMSO concentration was maintained at 0.5% (v/v) in all conditions. Each concentration was tested in triplicate in three independent experiments.

PrestoBlue™ Cell Viability Reagent (Thermo Fisher Scientific) was added directly to the culture medium at 10% (v/v), and plates were incubated for 3 h at 37°C. Fluorescence was measured using a Tecan Spark multimode plate reader (λ_ex_ = 531 nm, λ_em_ = 572 nm). Cell viability was expressed as a percentage relative to untreated control cells. EC₅₀ values were determined by nonlinear regression using GraFit 5.0 (Erithacus Software, UK).

### ALDH activity measurements in cell extracts

For determination of ALDH activity in cellular extracts, A549 and A172 cells were harvested during exponential growth phase and pelleted by centrifugation. Cell pellets were lysed using M-PER™ Mammalian Protein Extraction Reagent (Thermo Fisher Scientific) according to the manufacturer’s instructions. Total protein concentration was determined by the Bradford method.

ALDH activity was measured fluorometrically by monitoring NADH formation as described above for recombinant enzymes. Assays were performed at 25°C in 50 mM HEPES buffer pH 8.0 containing 50 mM MgCl₂, using saturating concentrations of hexanal and NAD⁺. For inhibition experiments, cell extracts were pre-incubated with increasing concentrations of Z34 for 20 min prior to activity measurements. IC₅₀ values were determined by non-linear regression analysis using GraFit 5.0 (Erithacus Software, UK) as described above.

## ACKNOWLEDGMENTS

This research was partially funded by the Spanish Ministerio de Ciencia, Innovación y Universidades (Agencia Estatal de Investigación, grant number PID2023-150696NB-I00 /MCIU/ AEI / 10.13039/501100011033 /FEDER, UE). This research was also supported by the Agència de Gestió d’Ajuts Universitaris i de Recerca (AGAUR, grup consolidat N-TETRASCAN 2021SGR00135). RP obtained financial support from the company Advanced BioDesign through a research contract agreement with Universitat Autònoma de Barcelona. We are indebted to Dr. Salvador Bartolomé and the Servei de Genòmica i Espectroscòpia de Biomolècules (SGiEB) for providing access to the Cary Eclipse fluorescence spectrometers (Agilent). We also thank MSc Oscar Palacios Ruiz and the Servei d’Anàlisi Química (SAQ) for providing access to the mass spectrometry facilities and for expert technicial assistance with sample preparation and ESI-MS sample injection. We thank Dr. Pablo Guerra from the IBMB-CSIC CryoEM Platform for assistance during the sample preparation and microscope data acquisition. The authors acknowledge funding from Project, IU16-014045 (CRYO-TEM) from Generalitat de Catalunya and by “ERDF A way of making Europe”, by the European Union. We also acknowledge Diamond Light Source for time on beamline Krios IV at the UK national electron Bio-Imaging Centre (eBIC) under proposal BI38805-6, and Yuriy Chaban for assistance during the data acquisition.

## Data availability

The structural data have been deposited in the Protein Data Bank (PDB code: 31DB) and are also reported in supplementary information.

## SUPPLEMENTARY MATERIAL

**Figure S1.**
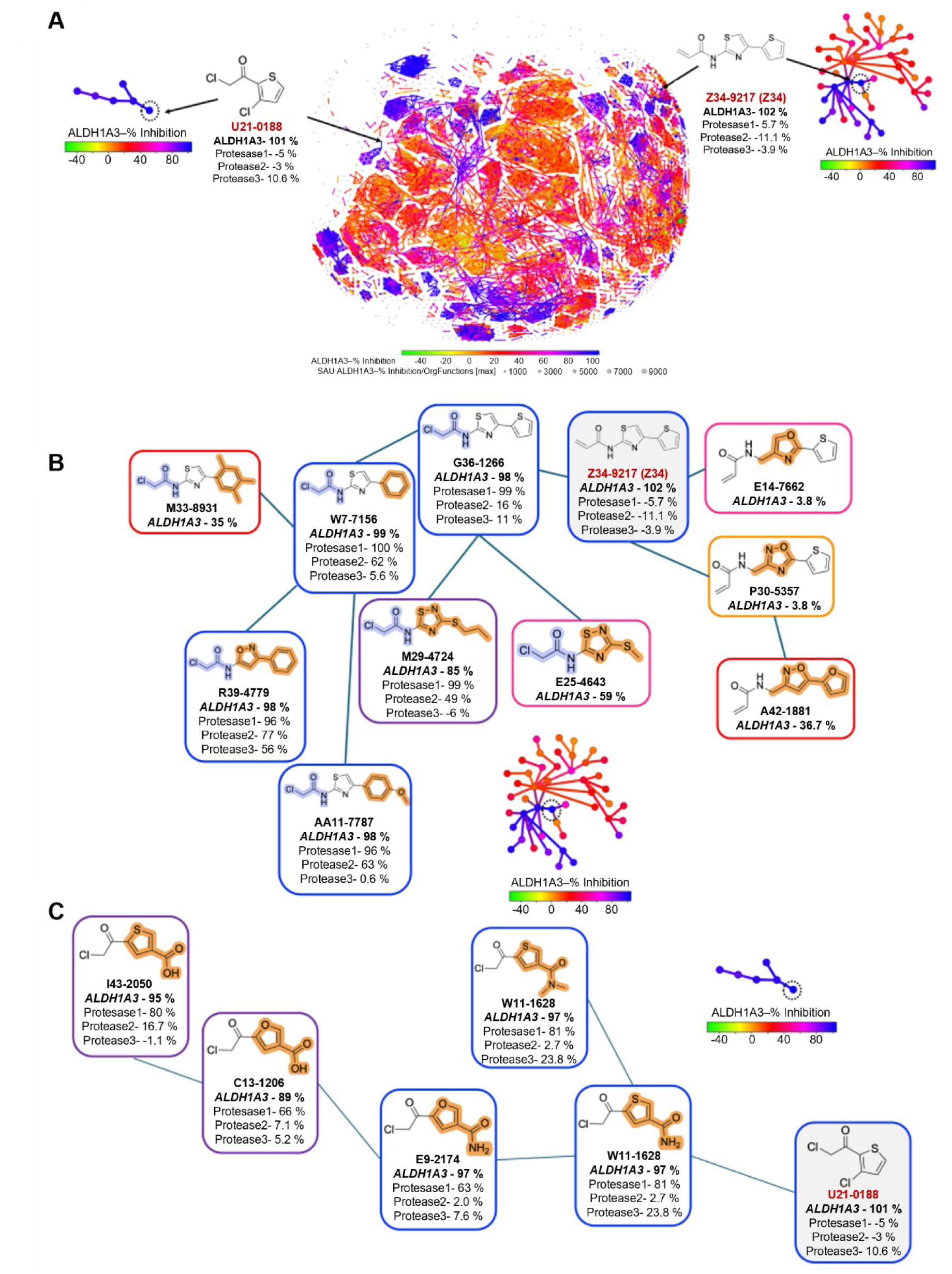
Structure-activity relationship analysis. **(A)** 2-D scatter plot showing structure activity landscape index (SALI) using OrgFunctions for the 3200 tested compounds. Each cluster shows structure similarity with variable activity. The locations of the two hit compounds with the neighborhood plot are marked by the arrow in the cluster. **(B-C)** The structure-activity relationship comparison with other active compounds in the neighborhood plot for Z34-9217 (Z34) (B) and U21-0188 (C).

**Figure S2.**
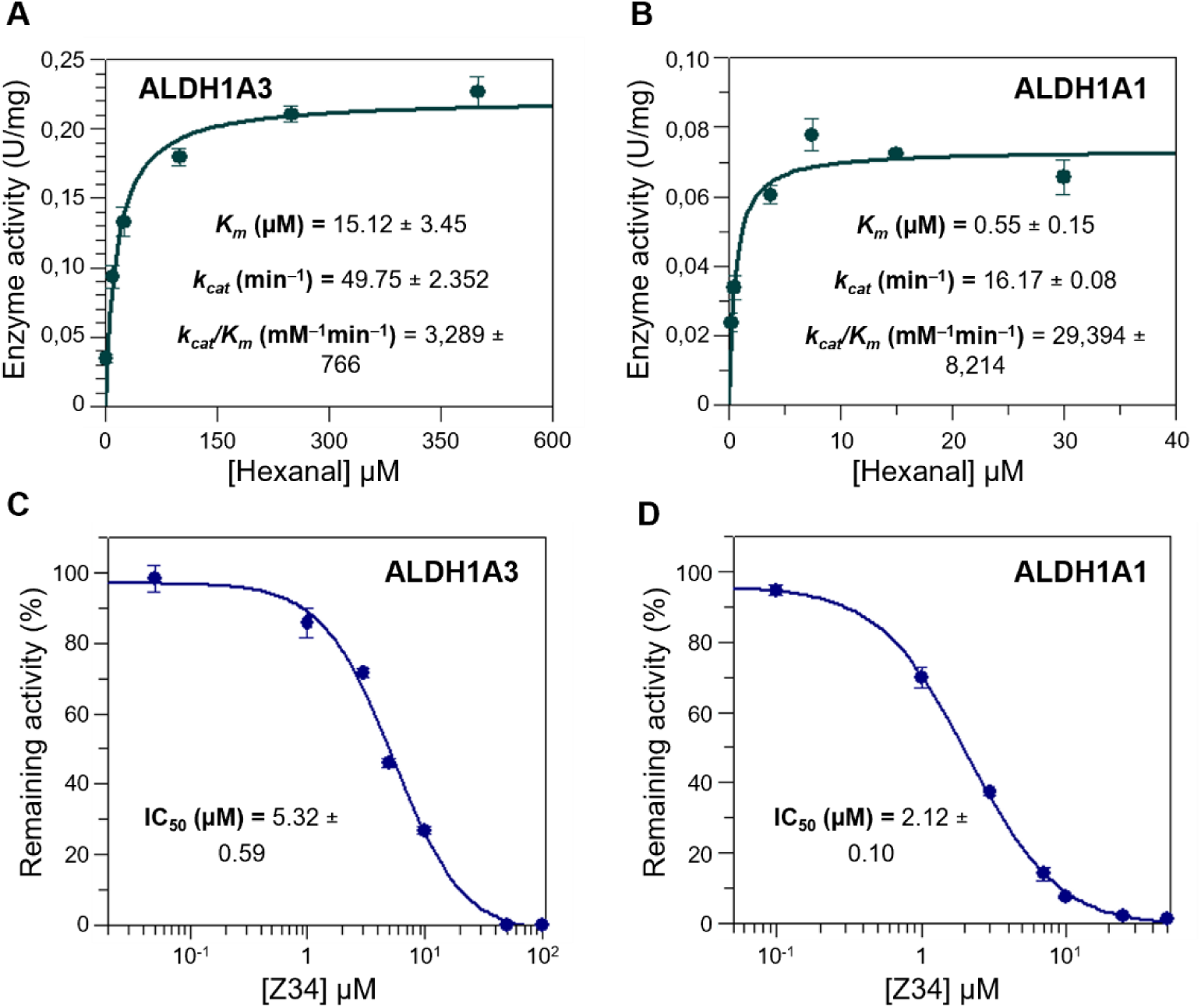
Kinetic characterization of ALDH1A3 and ALDH1A1. **(A)** Kinetic constants of ALDH1A3 with hexanal as a substrate. Enzymatic activity was measured fluorometrically in 50 mM HEPES, 50 mM MgCl₂, pH 8.0 at 25 °C. *k_cat_* value was calculated using a molecular weight of 224,000. Values are presented as mean ± standard error from triplicate measurements. **(B)** Kinetic constants of ALDH1A1 with hexanal as a substrate. Enzymatic activity was measured fluorometrically in 50 mM HEPES, 0.5 mM EDTA, pH 8.0 at 25 °C. *k_cat_* value was calculated using a molecular weight of 220,000. Values are mean ± standard error from triplicate measurements. **(C)** Sigmoidal representation used to determine the IC_50_ of Z34 for ALDH1A3 after 20 min pre-incubation in 50 mM HEPES, 50 mM MgCl₂, pH 8.0 at 25 °C. Values are mean ± standard deviation from duplicate measurements, and the IC_50_ is presented as mean ± standard error. **(D)** Sigmoidal representation used to determine the IC_50_ of Z34 for ALDH1A1 after 20 min pre-incubation in the buffer conditions described in panel B.

**Figure S3.**
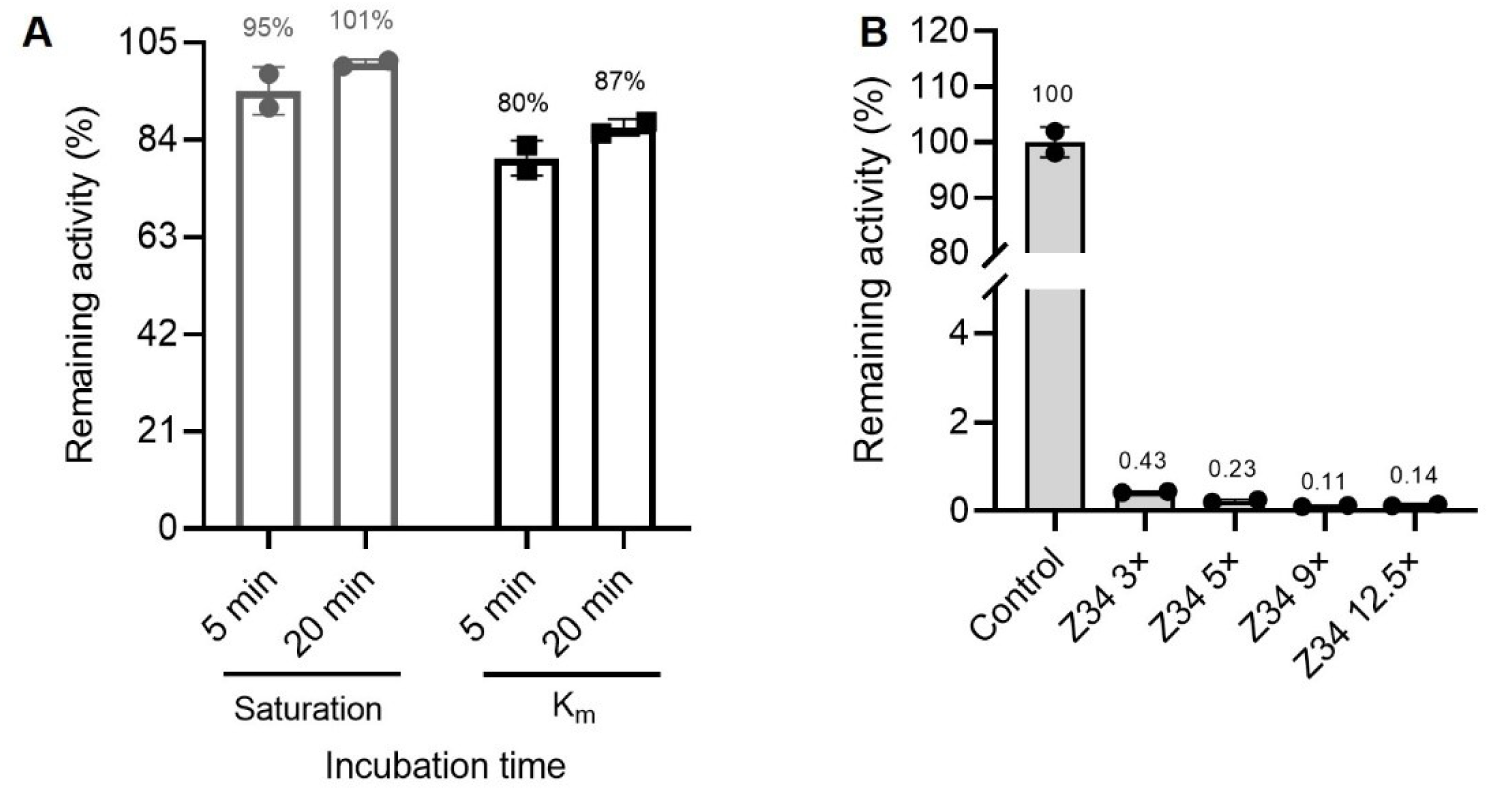
Assay validation and control experiments. **(A)** Initial inhibition screening of ALDH1A3 using 10 µM Z34 under assay conditions containing 50 mM HEPES, 50 mM MgCl₂, 5 mM DTT, pH 8.0 at 25 °C. **(B)** Enzymatic activity of ALDH1A3 following overnight incubation with Z34 at increasing molar equivalents relative to the monomer (3×, 5×, 9×, and 12.5×). Protein samples (8.4 µM, calculated based on the tetrameric molecular weight) were incubated at 25 °C in the presence of 4.2% (v/v) DMSO (control) or the indicated Z34 concentrations. Remaining activity was measured in 50 mM HEPES buffer containing 50 mM MgCl₂ (pH 8.0) at 25 °C using 250 µM hexanal as a substrate.

**Figure S4.**
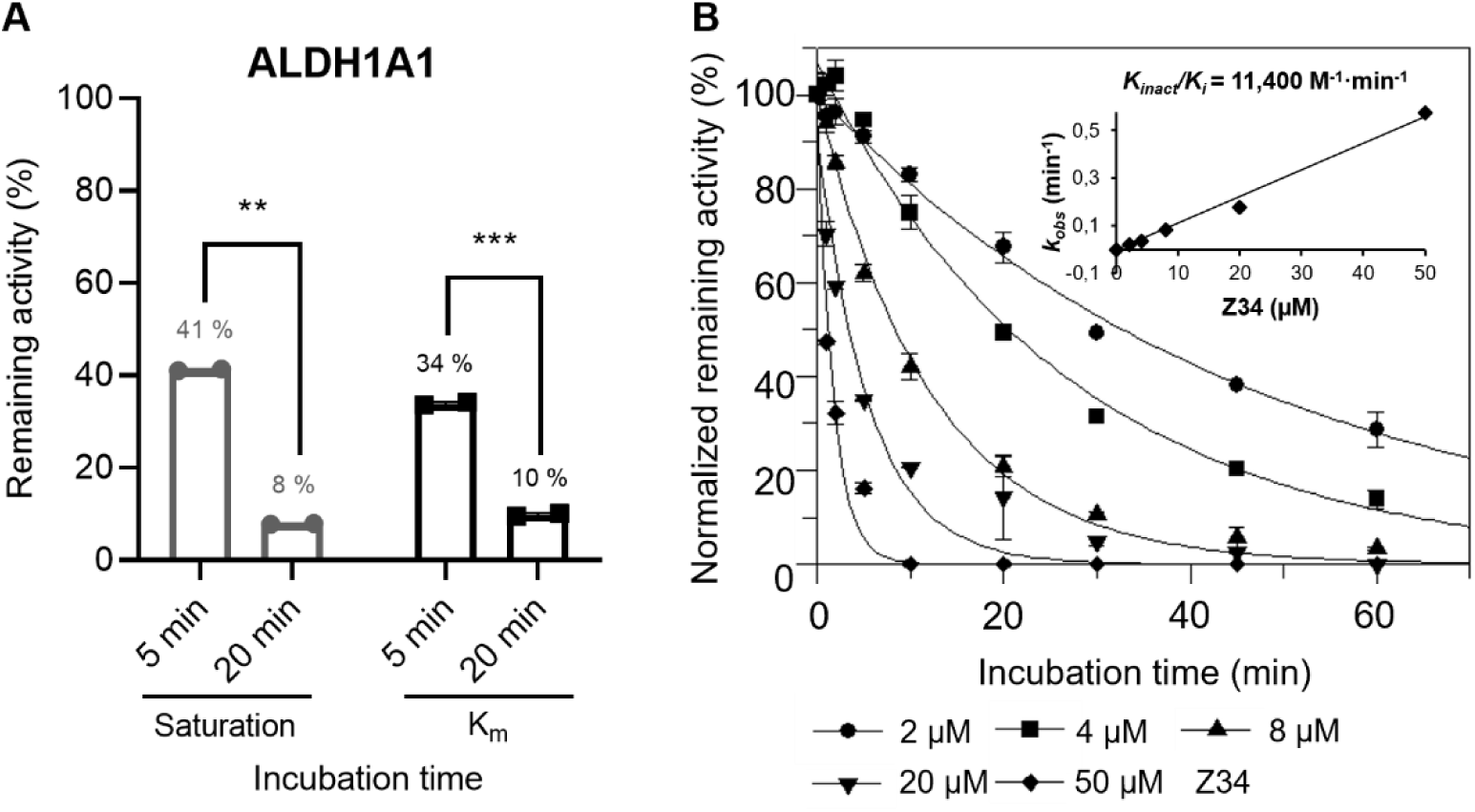
Biochemical characterization of ALDH1A1 inhibition by Z34. **(A)** Initial inhibition screening of ALDH1A1 using 10 µM Z34 at saturating (30 µM hexanal) or near-*K_m_* (0.5 µM hexanal) substrate concentrations after 5 and 20 min pre-incubation. Reactions were performed in 50 mM HEPES buffer containing 0.5 mM EDTA (pH 8.0) at 25 °C. Statistical analysis was performed using one-way ANOVA with multiple comparisons. **(B)** Determination of apparent inactivation rate constants (*k_obs_*) for ALDH1A1 at increasing Z34 concentrations (1×, 2×, 4×, 10×, and 20× IC_50_) using pre-incubation times ranging from 0 to 60 min. Dependence of *k_obs_* on inhibitor concentration was used to calculate *k_inact_*/*K_i_*. Experiments were performed in duplicate, and kinetic parameters were obtained by non-linear regression using GraFit 5.0 (Erithacus Software).

**Figure S5.**
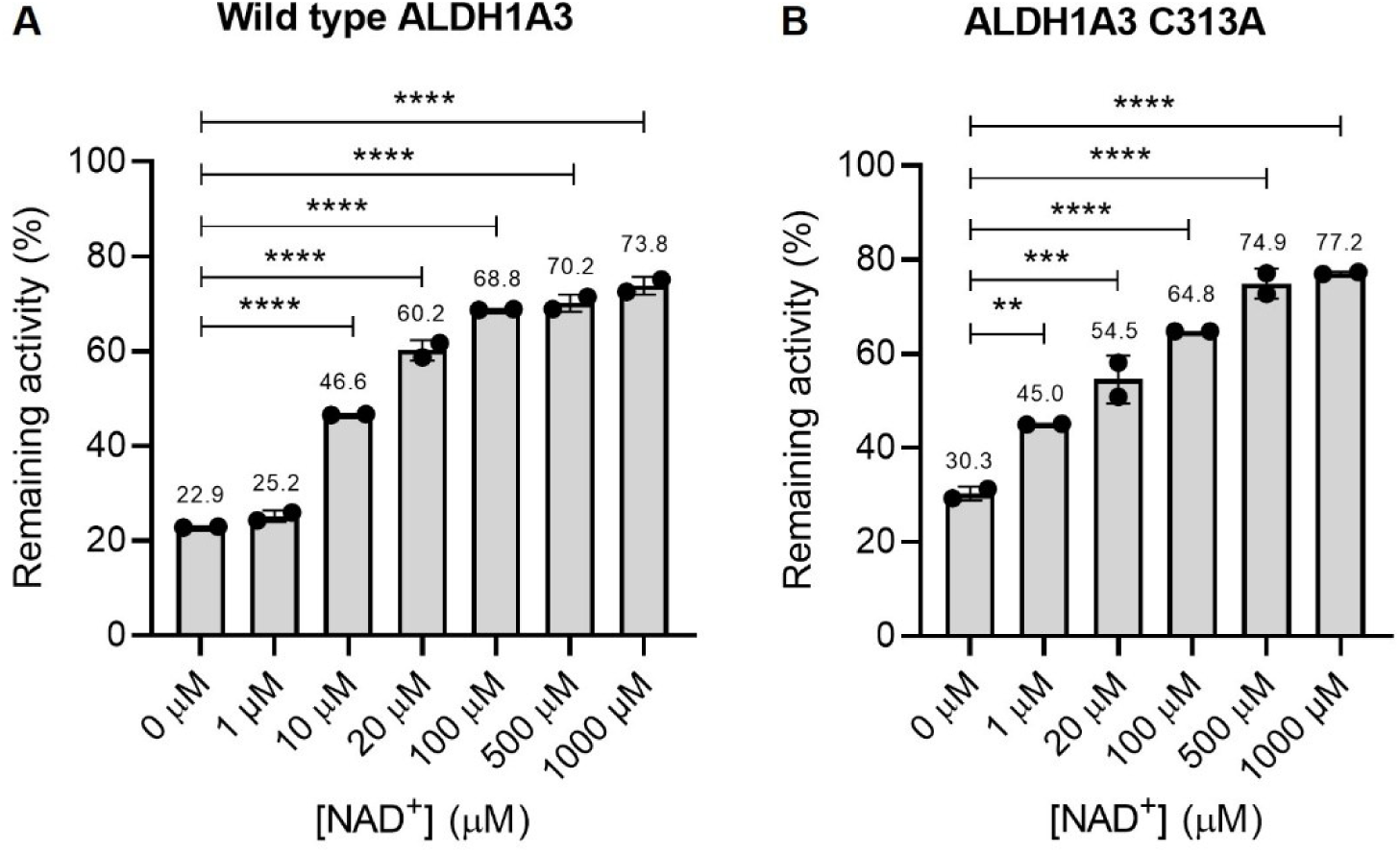
Additional analysis of NAD⁺ competition. **(A)** Bar chart showing ALDH1A3 enzymatic activity after 20 min incubation with 10 µM Z34 in the presence of NAD⁺ at concentrations ranging from 0 to 1000 µM. Values are presented as mean ± SD of duplicate measurements. **(B)** Enzymatic activity of the ALDH1A3 C313A mutant after 20 min incubation with 10 µM Z34 in the presence of increasing NAD⁺ concentrations (0–1000 µM). Values are presented as the mean ± SD of duplicate measurements.

**Figure S6.**
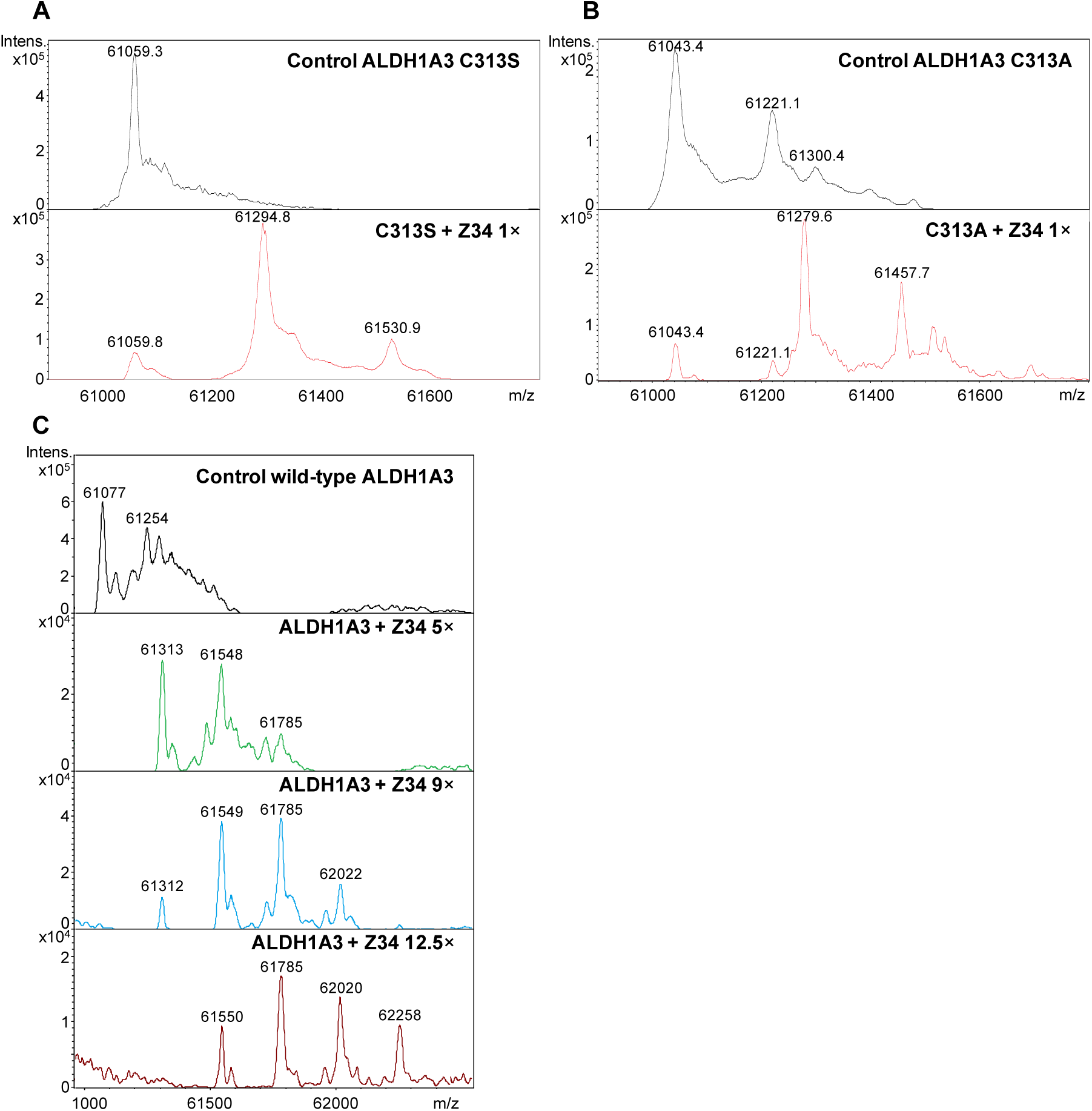
Additional mass spectrometry analysis of Z34 interaction with ALDH1A3. **(A)** ESI-TOF mass spectrometry analysis of the ALDH1A3 C313S mutant after incubation with Z34. Control samples contained 1% DMSO, and treated samples correspond to 1× Z34 per monomer. Mass spectra show the monomeric protein species under each condition. **(B)** ESI-TOF mass spectrometry analysis of the ALDH1A3 C313A mutant under the same conditions described in panel A (control and 1× Z34 per monomer). **(C)** ESI-TOF mass spectrometry analysis of wild-type ALDH1A3 incubated with higher Z34 ratios (5×, 9×, and 12.5× equivalents per monomer). Additional mass shifts corresponding to multiple ligand-associated species were detected at elevated ligand concentrations.

**Figure S7.**
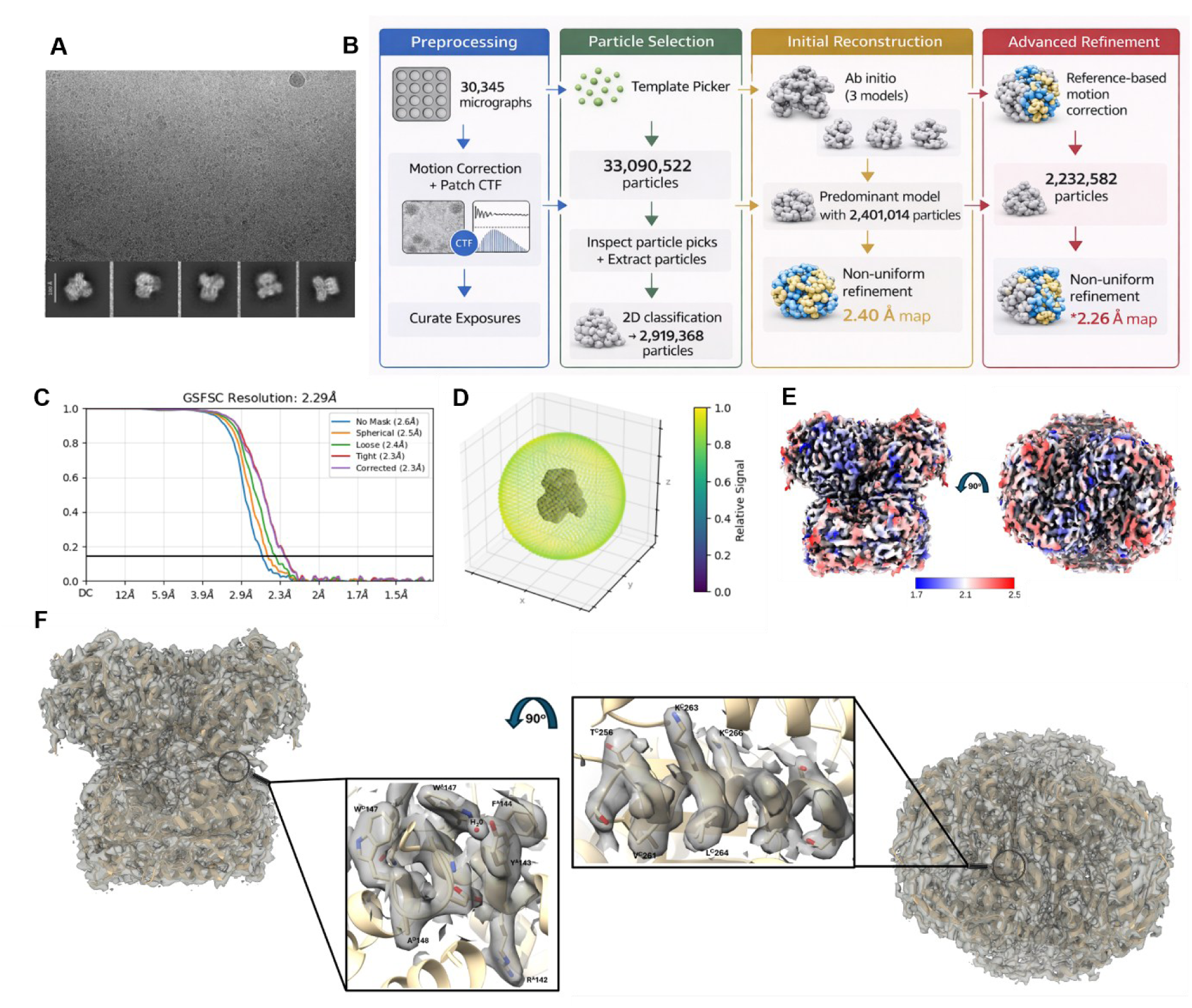
Cryo-EM data processing and structural reconstruction of ALDH1A3. **(A)** Representative cryo-EM micrograph of vitrified ALDH1A3 particles together with selected 2D class averages. **(B)** Schematic overview of the cryo-EM data processing workflow, including particle preprocessing, particle selection, initial 3D reconstruction and subsequent refinement steps. **(C)** Gold-standard Fourier shell correlation (FSC) curves used to estimate the global resolution of the reconstruction. **(D)** Spatial distribution of images used to obtain the map of ALDH1A3. **(E)** Cryo-EM density map colored according to local resolution. **(F)** The cryo-EM density map with the final atomic model of ALDH1A3. A close-up view highlights the quality of the map around representative side chains to illustrate the level of structural details available.

**Figure S8.**
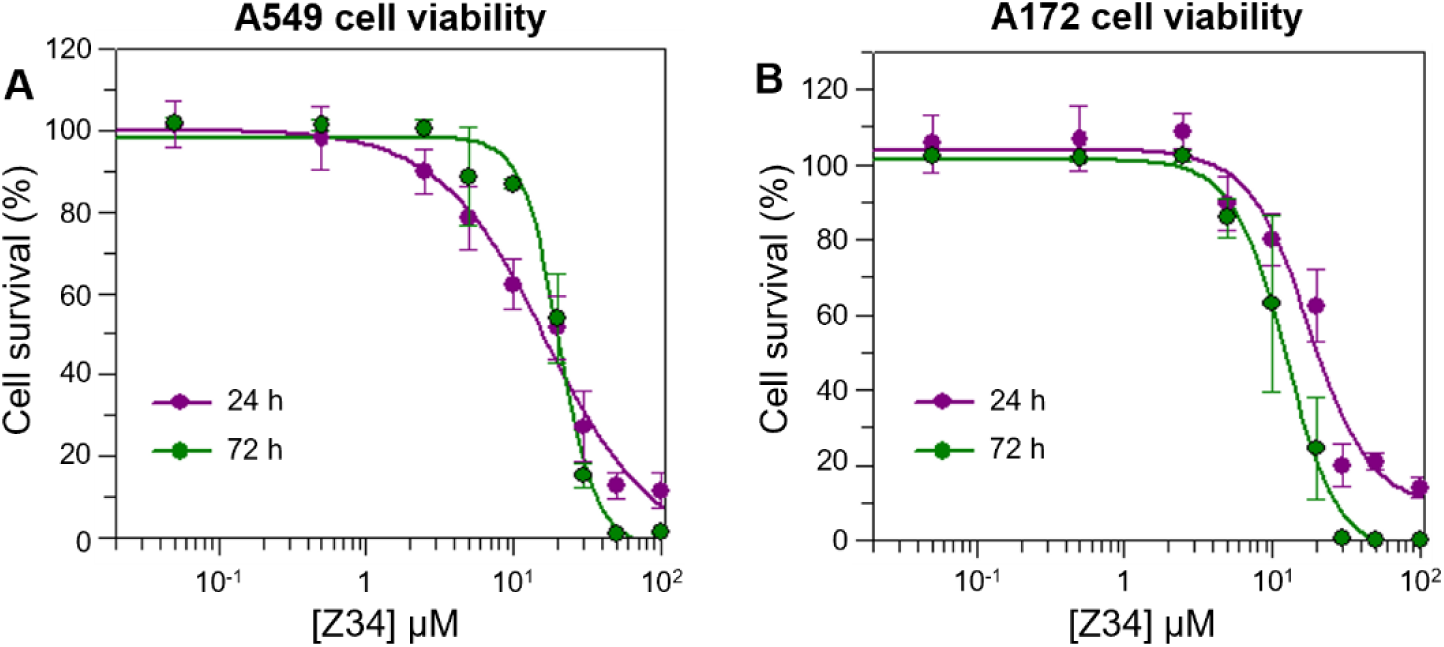
Time-dependent effects of Z34 on cancer cell viability. **(A)** Effect of Z34 on A549 cell viability after 24 and 72 h treatment. Dose–response curves were used to determine EC₅₀ values at each time point. EC₅₀ values were 16.6 ± 4.3 µM and 20.3 ± 1.3 µM after 24 and 72 h treatment, respectively. Data represent mean ± SD from three independent experiments, each performed with three technical replicates per condition. **(B)** Effect of Z34 on A172 cell viability after 24 and 72 h treatment. Dose–response curves were used to determine EC₅₀ values at each time point. EC₅₀ values were 17.7 ± 3.2 µM and 12.3 ± 0.8 µM after 24 and 72 h treatment, respectively. Data represent mean ± SD from three independent experiments, each performed with three technical replicates per condition.

**Table S1.**
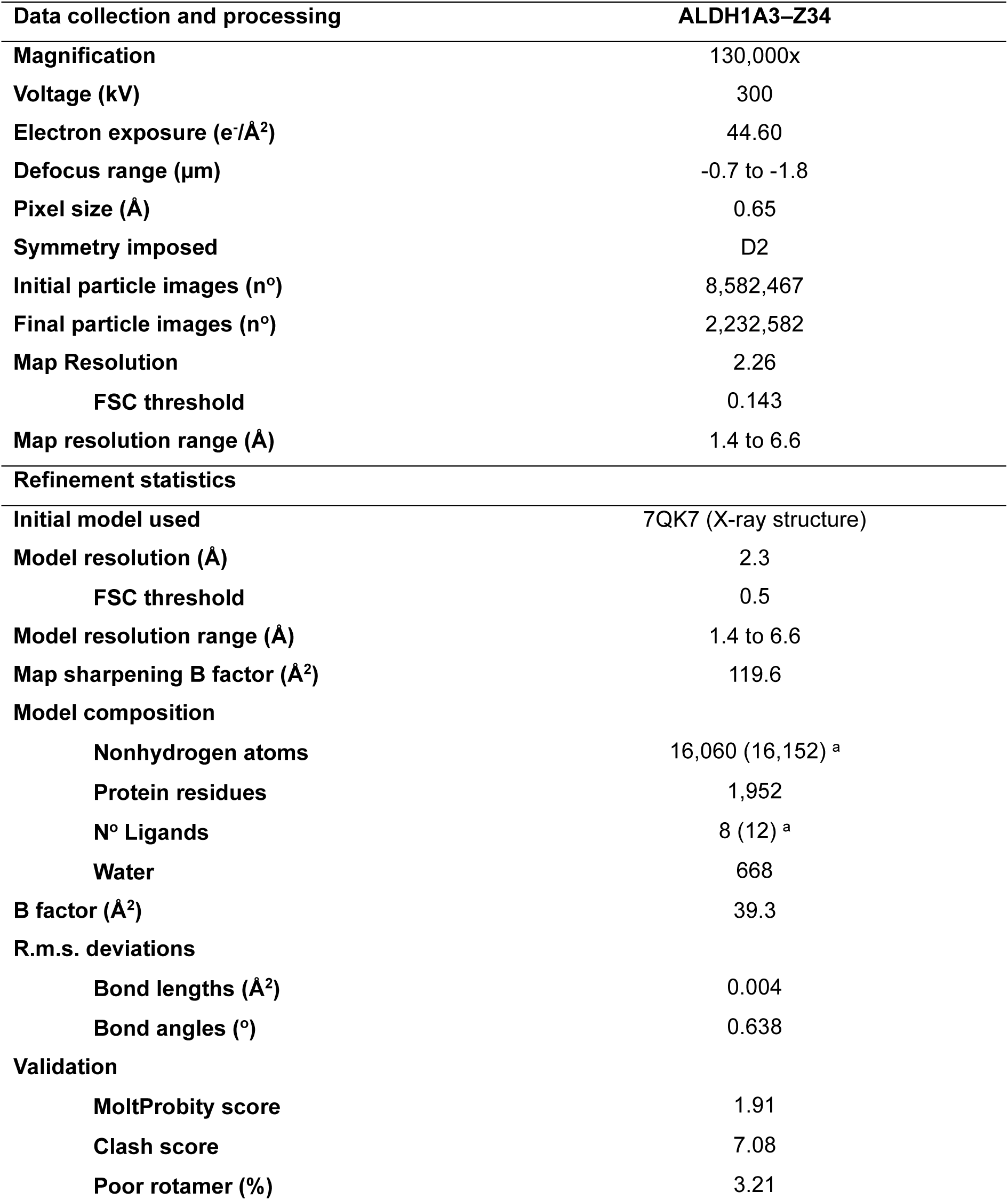
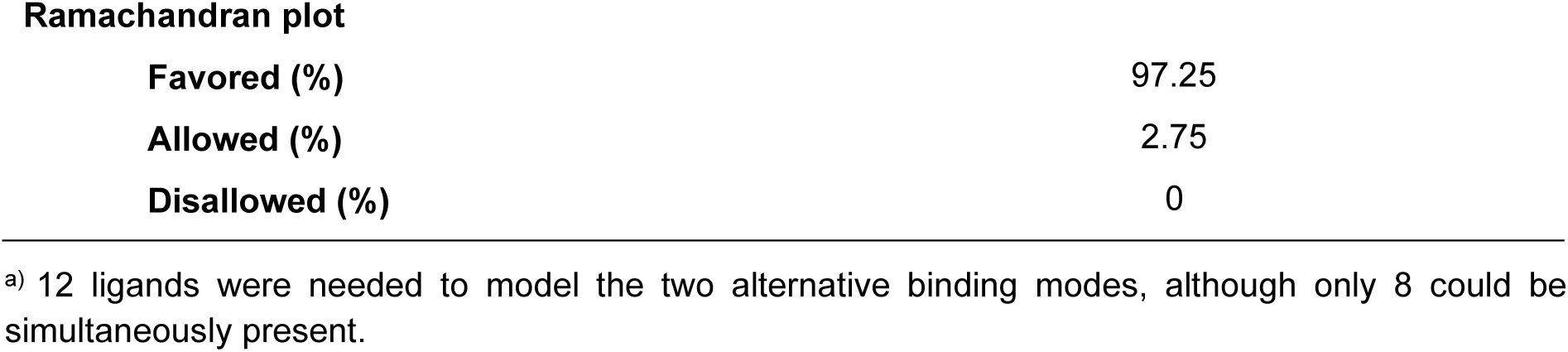
Cryo-EM data collection, processing and refinement statistics for the ALDH-Z34 complex.

| Data collection and processing | ALDH1A3–Z34 |
| --- | --- |
| Magnification | 130,000x |
| Voltage (kV) | 300 |
| Electron exposure (e<sup>-</sup>/Å<sup>2</sup>) | 44.60 |
| Defocus range (μm) | -0.7 to -1.8 |
| Pixel size (Å) | 0.65 |
| Symmetry imposed | D2 |
| Initial particle images (n°) | 8,582,467 |
| Final particle images (n°) | 2,232,582 |
| Map Resolution | 2.26 |
| FSC threshold | 0.143 |
| Map resolution range (Å) | 1.4 to 6.6 |
| Refinement statistics |  |
| Initial model used | 7QK7 (X-ray structure) |
| Model resolution (Å) | 2.3 |
| FSC threshold | 0.5 |
| Model resolution range (Å) | 1.4 to 6.6 |
| Map sharpening B factor (Å<sup>2</sup>) | 119.6 |
| Model composition |  |
| Nonhydrogen atoms | 16,060 (16,152) <sup>a</sup> |
| Protein residues | 1,952 |
| N° Ligands | 8 (12) <sup>a</sup> |
| Water | 668 |
| B factor (Å<sup>2</sup>) | 39.3 |
| R.m.s. deviations |  |
| Bond lengths (Å<sup>2</sup>) | 0.004 |
| Bond angles (°) | 0.638 |
| Validation |  |
| MoltProbity score | 1.91 |
| Clash score | 7.08 |
| Poor rotamer (%) | 3.21 |
| **Ramachandran plot** |  |
| **Favored (%)** | 97.25 |
| **Allowed (%)** | 2.75 |
| **Disallowed (%)** | 0 |

<sup>a)</sup> 12 ligands were needed to model the two alternative binding modes, although only 8 could be simultaneously present.

**Table S2.**
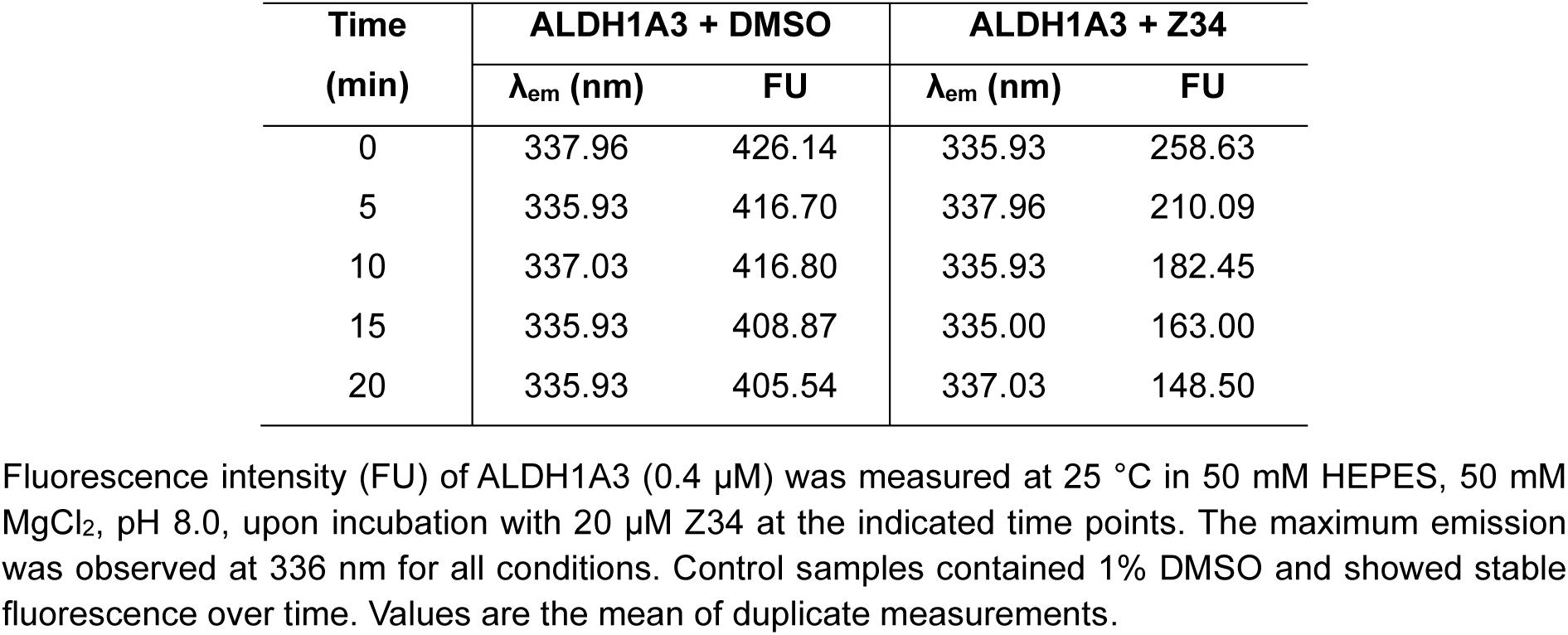
Intrinsic tryptophan fluorescence of ALDH1A3 upon incubation with Z34.

